# Leveraging state-of-the-art AI structure predictors to generate atomic resolution ensembles of intrinsically disordered proteins

**DOI:** 10.64898/2026.11.10.765624

**Authors:** Vincent Schnapka, Nicolas Portal, Massimiliano Bonomi

## Abstract

Intrinsically disordered proteins (IDPs) are central to many biological processes, but their function depends on dynamic conformational ensembles rather than a single structure. Despite the impact of artificial intelligence on protein structure prediction, determining these ensembles at atomistic resolution remains challenging. Recently, we introduced bAIes, a framework that leverages the structural information encoded in AlphaFold2 distograms to generate conformational ensembles of IDPs at atomistic resolution. Here, we extend this framework to six state-of-the-art structure prediction tools (AlphaFold2, AlphaFold3, Boltz-1, Boltz-2, ESMFold, and ESMFold2) and systematically compare the structural information encoded in their predicted inter-residue distance distributions. Despite differences between predictors, we show that bAIes generates ensembles with comparable accuracy against experimental NMR and SAXS data, and competitive performance relative to state-of-the-art all-atom simulations and alternative IDP ensemble generation methods. These results demonstrate that bAIes is robust to the choice of structure predictor and that the distance distributions generated by different deep learning architectures contain valuable information about the conformational preferences of disordered proteins. More broadly, our work highlights the potential of combining AI-derived structural information with physics-based modeling to generate atomistic ensembles of intrinsically disordered proteins and regions.

**Visual abstract:** 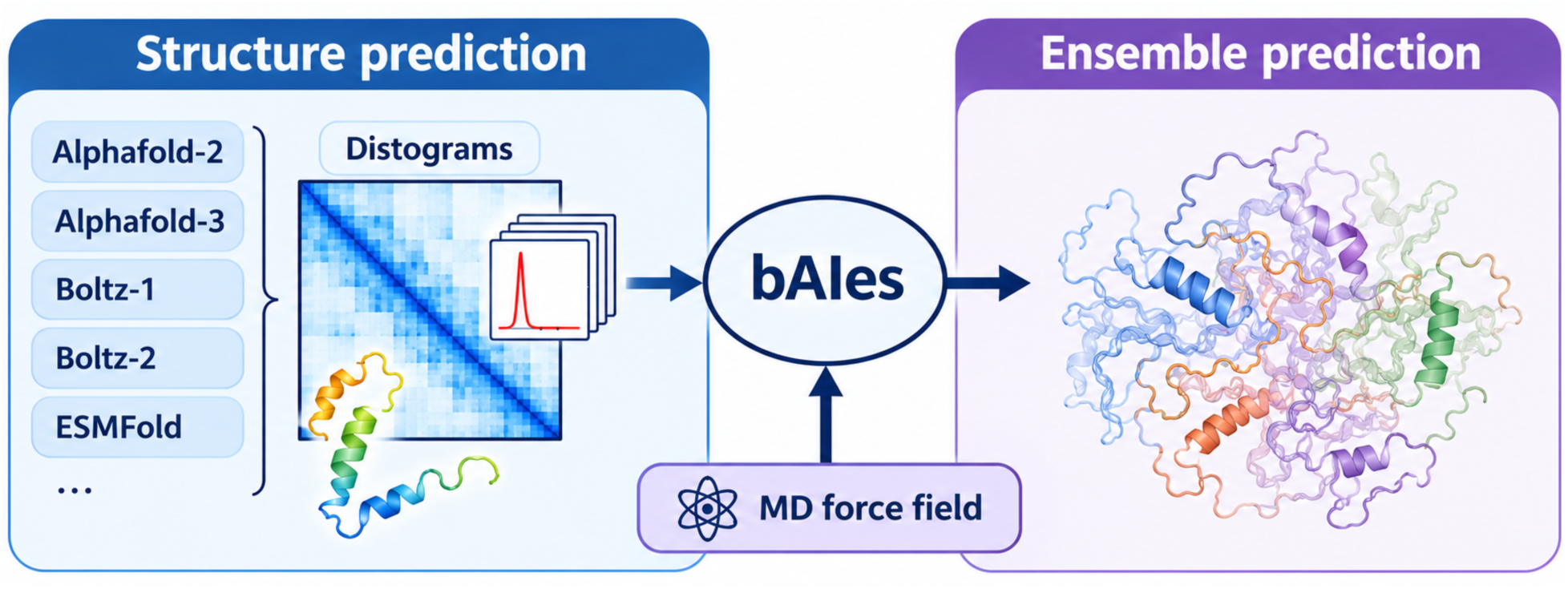

## Introduction

Intrinsically disordered proteins (IDPs) represent approximately one-third of the human pro-teome^1^ and play fundamental roles in numerous biological processes.^2^ They are characterized by the absence of a stable three-dimensional structure, making them challenging to characterize using a single experimental technique. Instead, IDPs are studied using complementary experimental techniques such as nuclear magnetic resonance (NMR) spectroscopy, small-angle X-ray scattering (SAXS), and Förster resonance energy transfer (FRET).^3,4^ Each of these techniques provides sparse and often ensemble-averaged structural information, making the experimental characterization of IDPs costly and the accumulation of structural ensemble data more challenging than for folded proteins.^5^ These limitations make the prediction of IDP conformational ensembles using computational methods particularly desirable yet challenging.

The prediction of IDP conformational ensembles has been approached using diverse *in silico* strategies.^6^ Recent developments in coarse-grained (CG) models for molecular dynamics (MD) simulations have enabled successful predictions of the global dimensions and phase behavior of IDPs,^7–12^ with studies extending to the entire human disordered proteome.^13^ In contrast, conventional all-atom MD approaches provide a detailed atomistic description but remain limited by computational costs and force field inaccuracies.^14–16^ Fragment-based approaches also remain popular, as they significantly reduce the computational cost of generating IDP ensembles by assembling conformations from peptide fragments derived from fragment modeling^17,18^ or statistical analyses of protein structure databases.^19–23^ More recently, several deep learning approaches for ensemble generation have been introduced.^24–30^ Most of these models are trained on data derived from MD simulations, fragment-based methods, and/or structure prediction tools, making their performance dependent on the scope and quality of their training datasets. In addition, they often generate either coarse-grained conformations that require reconstruction at the atomistic level or some nonphysical structures that must be filtered out in a subsequent step.

Recently, we introduced an alternative strategy through bAIes, an MD-based tool that integrates information from AlphaFold2 (AF2) to generate IDP conformational ensembles at atomistic resolution.^31^ We showed that bAIes competes with state-of-the-art all-atom MD methods in reproducing various types of experimental data, while requiring only a fraction of the computational cost. This study demonstrated that the structural information encoded in AF2 predictions is valuable for conformational ensemble prediction. However, since the introduction of AF2, numerous other structure prediction tools have emerged, featuring different architectures and training datasets, raising the question of whether they could also be leveraged for IDP ensemble prediction with bAIes.

AlphaFold3 (AF3)^32^ and Boltz-1^33^ expand the scope of structure prediction to other biomolecules and introduce diffusion-based generative modules for structure generation, replacing the deterministic structure module of AF2. Boltz-2^34^ builds on Boltz-1’s architecture but expands its training set beyond static structures to include structural ensembles derived from NMR and molecular dynamics trajectories, exposing the model to local fluctuations and global conformational heterogeneity. ESMFold,^35^ by contrast, leverages the ESM-2 protein language model to predict structures directly from sequence, without requiring a multiple sequence alignment (MSA). ESMFold2^36^ builds on the newer ESMC language model family, combining its representations into a pair representation that is iteratively refined and decoded into atomistic structures via a diffusion module, reportedly achieving higher accuracy and throughput than its predecessor.

Despite these differences in architecture and training strategy, all these tools produce distograms, three-dimensional arrays encoding predicted pairwise distance distributions between residues in a protein sequence. These distograms were used by bAIes with AF2 to generate IDP conformational ensembles in our previous work,^31^ have been shown to be useful for *in silico* drug design,^37^ and provide a suitable protein representation for machine learning.^38^ The emergence of these new structure prediction methods raises several questions: how do the distograms differ across predictors? How does the accuracy of the ensembles generated with bAIes vary across predictors? Do some predictors provide more informative distograms than AF2? And how does bAIes compare with recent deep learning approaches for ensemble generation?

Here, we extend bAIes to six AI structure predictors (AF2, AF3, Boltz-1, Boltz-2, ESMFold, and ESMFold2) and provide a new implementation that scales efficiently to long chains on GPUs. Using datasets of well-characterized IDPs with experimental NMR and SAXS data,^28,31^ we benchmark bAIes against all-atom MD simulations and state-of-the-art IDP ensemble generators. All tested predictors can be used with bAIes to generate atomistic IDP ensembles, with an average accuracy comparable to that of the original AF2-based implementation. bAIes captures local conformational properties across predictors, and our analysis identifies limitations in describing long-range and condition-dependent behavior. Overall, this work establishes bAIes as a flexible framework that combines molecular force fields with AI-derived structural information to generate IDP ensembles.

## Methods

### Overview of the bAIes approach

bAIes^31^ is based on a Bayesian inference framework that estimates the posterior probability *p*(*X|D*) of a biomolecular conformation *X* given prior physico-chemical knowledge and newly acquired data *D* as:

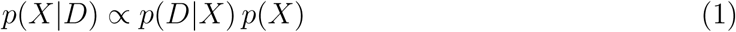

where *p*(*D|X*) is the likelihood function representing the probability of observing data *D* given the conformation *X*, and *p*(*X*) is the prior probability of *X* given existing physicochemical knowledge.

### The structural prior

In structural biology applications, the prior *p*(*X*) is often defined as:

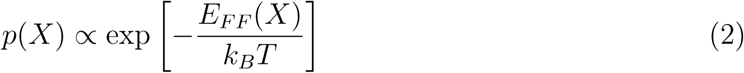

where *E_F F_* is a molecular mechanics force field encoding physico-chemical knowledge about the system. For bAIes, we developed a computationally efficient random coil model starting from the amber99SB-ILDN force field. ^39^ This force field was simplified by removing all non-bonded interactions except the repulsive Van der Waals terms and by correcting the backbone dihedral angle Φ and Ψ distributions with residue type-specific correction MAP terms.^40^ The resulting force field allows for rapid conformational sampling while yielding an accurate representation of local properties of random coil IDPs without structural bias.^31^

### The bAIes data

In bAIes, *D* represents the structural information provided by AI structure predictors, such as AF2, AF3, Boltz-1, and ESMFold. More specifically, this information is derived from a selection of pairwise distance distributions (distograms) between *C_β_* atoms (or *C_α_* for glycines) generated by the AI structure predictor. For each residue pair, the distogram is selected only if the two amino-acids are separated by at least 4 residues in sequence and if the distance corresponding to peak of the distogram is below a threshold corresponding to a typical contact distance for each type of amino-acid pair.^41^

### The data likelihood

For each selected pair of residues, we defined a Gaussian data likelihood:

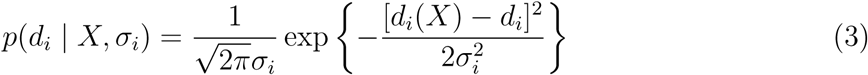

where *d_i_*(*X*) is the distance between a pair of *C_β_* atoms (*C_α_* for glycines) in conformation *X*, *d_i_* is the peak of a Gaussian function that best fits the distogram, and *σ_i_* is an uncertainty parameter that controls the spread of the distribution. Assuming that the distances are independent, we express the likelihood function for a set of *N* distances *D* = *{d_i_}* as:

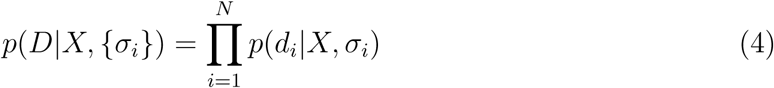

### The uncertainty prior

In bAIes, we treat the uncertainty parameters *σ_i_* as unknown variables and use a non-informative Jeffreys prior:

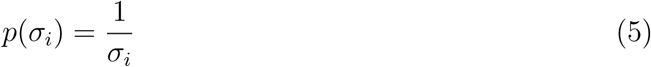

For each selected residue pair, this function is defined in the interval [*σ^min^,* +*∞*), where *σ^min^* is obtained from the Gaussian fit of the corresponding distogram. To avoid sampling the *σ_i_* parameters, the likelihood functions are marginalized along with their corresponding uncertainty priors, resulting in the following marginal data likelihoods:

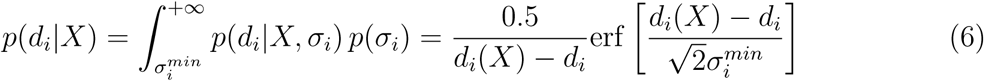

### The bAIes energy function

The final bAIes posterior distribution can be written as:

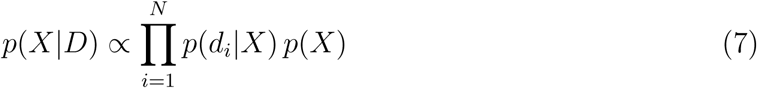

which can be sampled by a Molecular Dynamics simulation driven by the associated bAIes energy function:

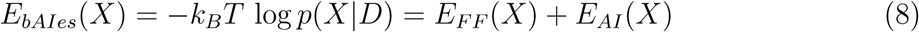

where *k_B_* is the Boltzmann constant, *T* the temperature of the system, *E_F F_* the random coil model, and:

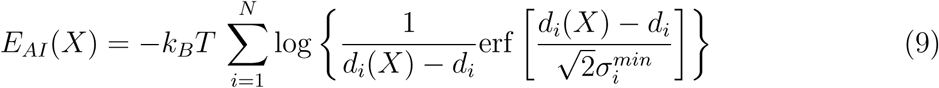

### Structure prediction and distogram extraction

The starting conformations and distograms for each bAIes simulation were generated on our local computing cluster using six structure prediction models, all run with default parameters: AlphaFold2^42^ (v2.2.0), AlphaFold3^32^ (v3.0.1), Boltz-1^33^ and Boltz-2^34^ (both run with the Boltz package, v2.1.1), ESMFold^35^ (*esmfold v1*, v2.0.1: https://github.com/facebookresearch/ESM), and ESMFold2^36^ (*esm*, v3.2.4: https://github.com/Biohub/esm). For AF2 and AF3, multiple sequence alignments (MSAs) were generated locally with each model’s built-in search pipeline and reference databases. For Boltz-1 and Boltz-2, which do not include a local MSA search pipeline, MSAs were retrieved from the public ColabFold MMseqs2 server (*–use msa server*). ESMFold and ESMFold2 predict structures directly from single sequences using protein language model embeddings and are run without MSAs.

For each model, distograms were extracted as pre-softmax logits of shape *N_res_ × N_res_ ×* 64, where *N_res_* denotes the number of protein residues. AF2 and ESMFold2 both allow the distograms to be obtained without modifying their code and were used as distributed. AF3, Boltz-1, Boltz-2, and ESMFold compute the distograms internally but do not return it, so their inference code was modified to save them. These modifications only expose an existing intermediate output and do not alter the predictions. Model weights and inference settings were otherwise left at their default values.

### Comparison of distograms across predictors using PeptoneDB-CS

PeptoneDB-CS is a curated set of 659 proteins with experimental NMR chemical shifts from the Biological Magnetic Resonance Data Bank (BMRB). ^28^ The dataset is balanced between predominantly ordered (329) and predominantly disordered (330) sequences, as defined by the mean per-residue G-score (0 = fully ordered, 1 = fully disordered; disordered if *>* 0.5).^28^ Structure predictions were performed to predict the structures and distograms of the 659 proteins of PeptoneDB-CS using each of the 6 structure predictors defined above, representing a total of 3954 structure predictions. Following our previous work,^31^ each pair of residues within the protein was considered in contact if: i) they are separated by at least 4 residues in the sequence, and ii) the most probable pairwise *C_β_ − C_β_* distance falls below a threshold^31,41^ corresponding to a typical contact distance for each type of amino-acid pair. The average number of inter-residue contacts was evaluated for each structure predictor and for the 329 ordered (g-score *<* 0.5) and 330 disordered proteins (g-score *>* 0.5) of the benchmark separately. The error bars were obtained from bootstrap analysis^43^ with 2000 resamples of the 329 or 330 independent data points (per-protein predicted distograms). The consensus matrices **M** in Figure S3 were obtained by quantifying and averaging over all of the *N* proteins (*N* =329 for ordered proteins and *N* =330 for disordered proteins), for each predictor pair (*A, B*), the proportion of inter-residue contacts predicted by predictor *A* also predicted by predictor *B*. The *M_A,B_* elements of each matrix were calculated as follows:

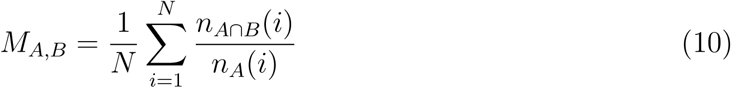

where *n_A_*(*i*) is the total number of contacts predicted by predictor *A* in protein *i*, and *n_A∩B_*(*i*) is the total number of contacts in common within the predictions of predictors *A* and *B* in protein *i*.

### bAIes simulation protocol

bAIes simulations were performed using OpenMM v. 7.6.0.^44^ While PLUMED^45^ was used in our previous work, ^31^ and despite the fact that PLUMED is compatible with OpenMM, we natively implemented bAIes in this software using a “CustomBondForce” in order to optimize computational efficiency. For each protein and structure predictor, bAIes simulations were prepared and carried out as follows:

- The top ranked predicted model was used as a starting point for the simulation, and the associated set of distograms was used for bAIes;
- The distograms were processed as described above to select the inter-residue contacts and extract the *d_i_* and *σ_i_* parameters with an in-house python script available on GitHub (https://github.com/COSBlab/bAIes-IDP);
- The predicted structure in PDB format was converted into an MD-ready PDB file with the pdb2gmx tool from GROMACS 2022.5^46^ to add the missing hydrogens;
- Energy minimization was performed using OpenMM’s L-BFGS local minimizer (”LocalEnergyMinimizer”) with default parameters;
- Production simulations were performed in the NVT ensemble at 298.1 K using a Langevin thermostat^47^ with a friction coefficient of 0.01 ps*^−^*^1^ and a time step equal to 2 fs. Bonds with hydrogen atoms were constrained using SHAKE.^48^

To construct the final bAIes ensembles, the protein coordinates were saved after each interval I in picoseconds defined as:

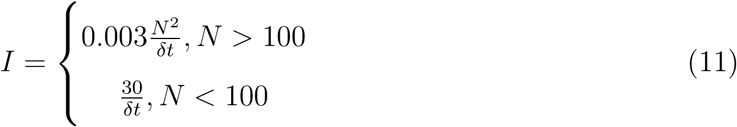

where *N* is the number of residues in the protein chain and *δt* is the simulation time step in ps. This formula ensured sufficient de-correlation of the saved frames, independently of the protein size as the simulation evolved.^9^ Simulation convergence with respect to the experimental quantities of interest was systematically assessed using blocking analysis, as described below.

### Computational performance across simulation platforms

To assess the computational performance of bAIes as a function of protein size, we constructed disordered protein chains composed of only glutamine residues (polyQ) with various sizes, from 10 to 500 residues, using PyMol.^49^ The computational efficiency of the coil model was then assessed for each polyQ protein with short 20,000-step simulations using either LAMMPS (2 Aug. 2023)^50^ or our novel OpenMM implementation. Simulation speed was evaluated over the production stage. For LAMMPS, we used the performance (ns/day) reported at the end of the production run. For OpenMM, we measured the elapsed wall-clock time of *simulation.step*() with the *time.perf counter*() command, followed by a state query (*getState*) to ensure GPU execution had completed, and converted it to ns/day using the integration timestep. LAMMPS simulations used one MPI task with eight OpenMP threads and a time step of 1 or 2 fs. OpenMM simulations used a 2 fs time step and ran either on eight CPU threads or with the CUDA platform on an NVIDIA A40 GPU. All CPU runs were performed on AMD EPYC 74F3 processors (24 cores).

The computational efficiency of bAIes was evaluated with OpenMM’s CUDA platform as a function of protein size and number of bAIes restraints on 2883 proteins from the DisProt database,^51^ a manually curated and annotated database of intrinsically disordered proteins and regions. bAIes simulations were performed for each protein using OpenMM as described above. One hundred frames were generated for each protein, and the interval after which protein coordinates were saved was defined in picoseconds as:

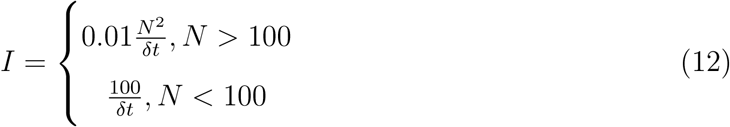

where *N* is the number of residues in the protein chain and *δt* is the simulation time step in ps. A subset of 2443 ensembles of the largest proteins with non-overlapping sequences, selected from the original set of 2883 proteins, was deposited in the Protein Ensemble Database (PED).^52^

### Back-calculation of NMR and SAXS observables

Chemical shift predictions were performed using SPARTA+^53^ and UCBShift.^54^ Both predictors were compared and yielded consistent results (Figure S11). ^3^*J_HNHA_* couplings were calculated using the Karplus relationship from Vogeli and Bax (7.97,-1.26 and 0.63 Hz).^55^ ^3^*J_CC_* couplings were calculated using a Karplus relationship from Li *et al.* (1.61,-0.93 and 0.55 Hz).^56^ Residual Dipolar Couplings (RDCs) were calculated using a local alignment window of 15 residues^57^ using PALES.^58^ The global alignment scaling factors were optimized to minimize the deviation between the calculated and experimental data points of equal sign.^31^ Paramagnetic relaxation enhancements were calculated as described in^31^ with a known rotameric ensemble of MTSL paramagnetic side chains ^59^ and a correlation time of 4 ns for the electron-nucleus interaction vector. Intensity ratios were calculated from Paramagnetic Relaxation Enhancement (PRE) using an INEPT time of 10 ms and an intrinsic transverse relaxation rate of 4 s*^−^*^1^. For the PeptoneDB-Integrative benchmark (Figure 5), intensity ratios were calculated following the PeptoneBench protocol of Invernizzi *et al.* ^28^ for consistency with the benchmark. Briefly, the PREs were calculated with DEERPREdict^60^ and converted to intensity ratios using an INEPT time of 10 ms and an intrinsic relaxation rate of 10 s*^−^*^1^ for HSQC-type data and 50 s*^−^*^1^ for the HMQC-type data (FLN5A3A3). Following this protocol, the correlation time for the electron-nucleus interaction vector was optimized by scanning values from 1 to 20 ns and keeping the value that minimizes the average Root Mean Square Error (RMSE) against experiments across each labeling site for each ensemble. All the SAXS profiles were calculated using Pepsi-SAXS^61^ using the same procedure. For each conformer, a theoretical profile is computed with a fixed scale factor of 1 and no offset; the hydration shell contrast *dρ* and effective atomic radius *r*_0_ were fixed at 3.34 *·* 10^−3^ *e*°*A*^−3^ and 1.68 °*A*, respectively.^62^ The ensemble-averaged profile was computed across all conformers, along with an optimal scale factor and offset to minimize the *χ*^2^ with respect to the experimental SAXS profile.

### Validation and benchmarking against all-atom simulations

To benchmark bAIes against all-atom MD simulations, we used the dataset defined in our previous work. ^31^ This dataset contains 17 IDPs equally divided into three categories: almost random coil IDPs (n=6), IDPs with residual structural elements (n=6), and IDPs for which we identified prediction errors with AF2 (n=5). Four of these IDPs are extensively characterized by NMR and SAXS as well as by long all-atom MD simulations (30 microseconds) performed with state-of-the-art force fields, which we used as a reference for comparison^14–16,63^ (Table S1).

The method validation and benchmark against all-atom MD simulations were performed as in our previous work. ^31^ For each IDP ensemble predictor model *m*, we defined the RMSE between calculated and experimental data points, *d_i_^calc^* and *d_i_^exp^* respectively, as:

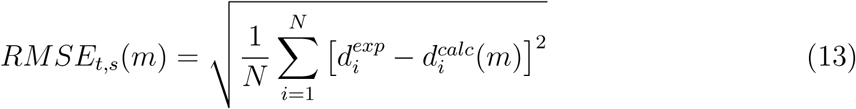

where *t* is the type of experiment (chemical shift, RDC, PRE, J-coupling, or SAXS) and *s* is a single experimental dataset (a labeling position for PREs, a specific nuclei for chemical shifts or a specific J-coupling between two nuclei). We defined the RMSE for a given experiment type as the average RMSE across the associated datasets:

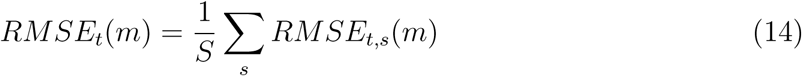

where *S* is the number of datasets associated with a given experiment type. RMSE values are expressed in the units of the corresponding experimental observable (ppm for chemical shifts, Hz for RDCs, and ^3^J-couplings; PRE intensity ratios and SAXS intensities in arbitrary units). Finally, we defined a global normalized RMSE (*RMSE_N_* (*m*)) relative to a reference model *m_ref_* (here the random coil prior) as:

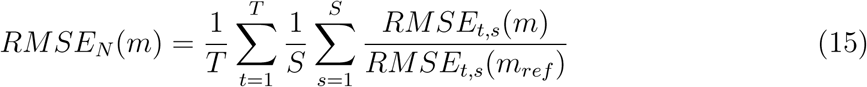

were *T* is the number of experimental data types (5 in our benchmark).

We calculated error bars using blocking analysis, where we divided each trajectory into three non overlapping blocks of equal length and computed the standard error of the mean of the calculated metrics across blocks.

### Benchmarking against other ensemble generation methods using PeptoneDB-Integrative

PeptoneDB-Integrative is a literature-curated set of 16 IDPs, each with several complementary experimental measurements, including NMR chemical shifts, backbone N-H RDCs, SAXS profiles, and PREs.^28^ For all these proteins, we compared the conformational ensembles generated with state-of-the-art approaches in Ref.^28^ and our bAIes ensembles with the experimental data in PeptoneDB-Integrative (Table S2). To do so, we followed the PeptoneBench protocol,^28^ with the exception that no reweighting against experimental data was applied. This choice reflects our aim to assess the raw predictive capability of each model, as required when no experimental data are available. Following PeptoneBench,^28^ each ensemble was evaluated against experiments as follows:

For chemical shifts and SAXS, a normalized RMSE (*nRMSE*) between calculated and experimental observables was calculated as follows:

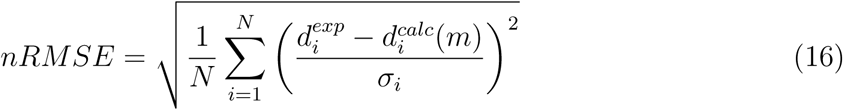

where *d_i_^calc^* and *d_i_^exp^* are the calculated and experimental data points, respectively, and *σ_i_* is a per-point uncertainty on the measured observable. This dimensionless quantity is interpretable: a value around 1 corresponds to an agreement within the estimated uncertainty. For SAXS, the experimental uncertainty was used for the *σ_i_*. For chemical shifts, the uncertainty was estimated as an interpolation between the original predictor uncertainties and the uncertainties estimated for random coil proteins:^64^

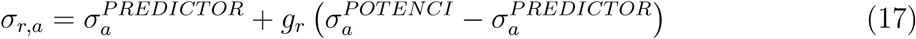

where *σ_a_^PREDICTOR^* is the UCBShift uncertainty associated with atom type *a*, and *σ_a_^POTENCI^* is the uncertainty associated with random-coil chemical shift predictions.^64^ *g_r_* is the per-residue disorder g-score that estimates the level of disorder in the considered protein from experimental secondary chemical shifts. This *g*-score-weighted uncertainty down-weights predictions on structured domains with respect to disordered regions, which tend to give much smaller deviations from experiments.

For RDCs, a per-protein Q-factor is calculated as an RMSE normalized by the root-mean-square of the experimental data points. It is calculated as follows:

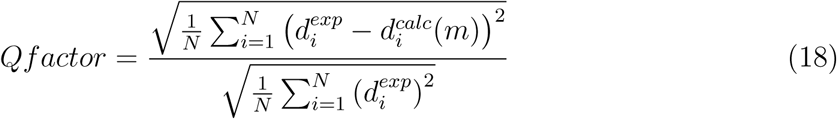

For PREs, agreement with the experiment was quantified by the RMSE computed on intensity ratios (*I_para_/I_dia_*). Experimental ratios above 1, which are nonphysical and attributable to noise, were set to 1.

## Results

### Consistency of IDP distograms across structure predictors

We first examined how the distograms produced by six AI structure predictors (AF2, AF3, Boltz-1, Boltz-2, ESMFold and ESMFold2) differ in the number of predicted inter-residue contacts and the overall shape of their distance distributions. As an example, the predicted models and distogram average distance matrices of PaaA2, a disordered protein with two important helical-propensity regions, are illustrated in Figure 1A. All methods tested systematically predict the two helices but differ in the identification of long-range contacts near the N-terminal domain. To obtain a broad, qualitative picture of how such differences depend on the structure predictor used, we analyzed 3954 predictions for a benchmark of 659 proteins with NMR backbone chemical shift data (PeptoneDB-CS^28^). The benchmark is perfectly balanced between ordered (mean g-score *<* 0.5; 329 proteins) and disordered (g-score *>* 0.5; 330 proteins) proteins (Figure S1).

**Figure 1:**
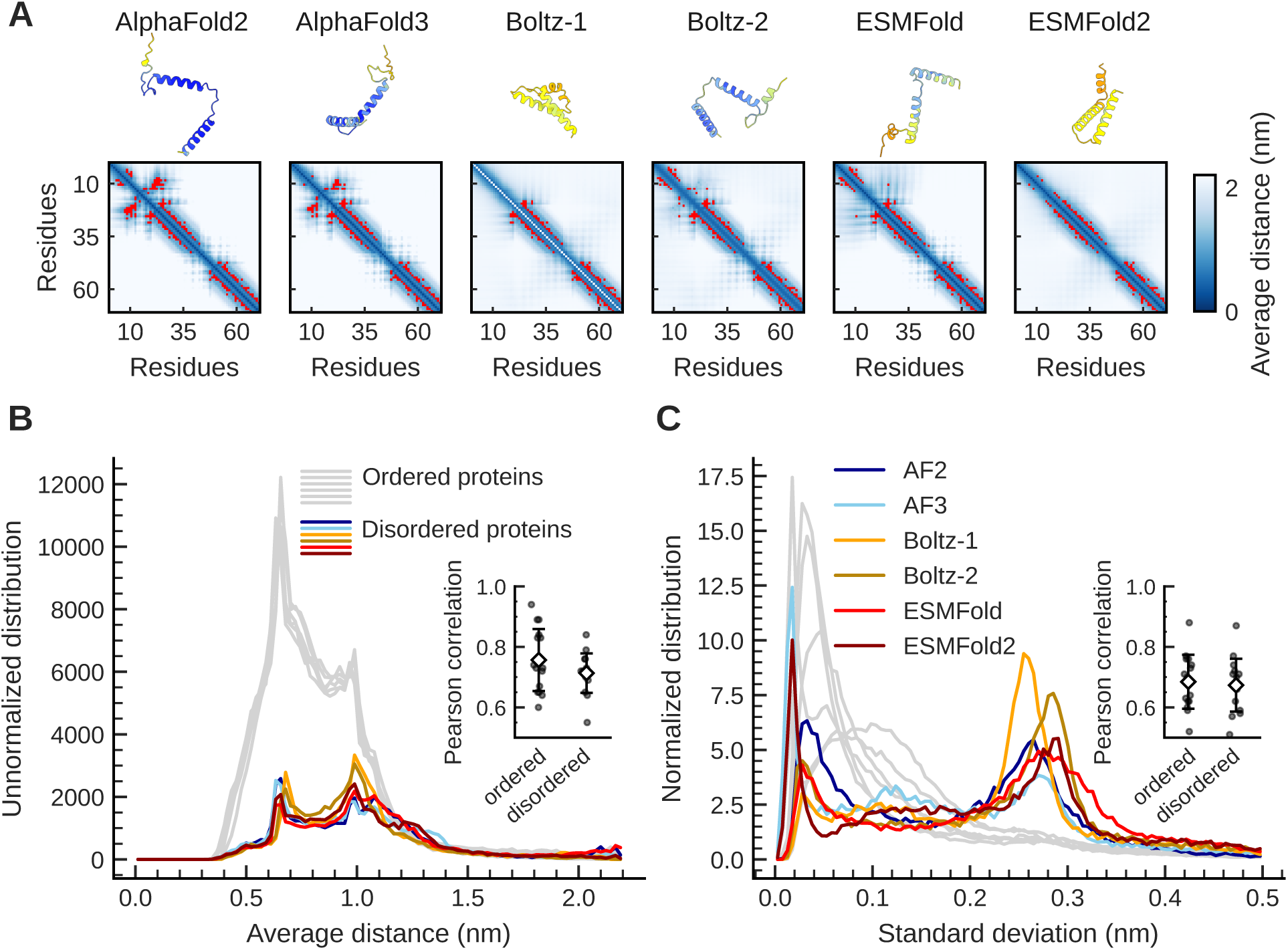
Comparison of distograms from state-of-the-art AI structure predictors. **A**: Structure predictions of PaaA2 from the six structure predictors, showing the predicted models (colored by pLDDT) and the corresponding pairwise average distance matrices. Residue-pair contacts, as defined in Methods, are highlighted in red. **B**: Distributions of distogram mean distances for the 659 proteins in the PeptoneDB-CS dataset, comprising 330 disordered and 329 ordered proteins. Predictions for disordered proteins are shown separately for AF2 (dark blue), AF3 (light blue), Boltz-1 (yellow), Boltz-2 (ochre), ESMFold (red), and ESMFold2 (dark red), while predictions for ordered proteins are shown in grey. Distributions are not normalized to highlight differences in the number of predicted contacts between ordered and disordered proteins. Inset: Mean Pearson correlation coefficients between predictors for disordered and ordered proteins. Black points represent individual predictor pairs, and error bars denote standard deviations across predictor pairs. **C**: As in B, but for distogram standard deviations. Distributions are normalized to highlight the larger fraction of high-spread distograms in disordered proteins.

On average across the entire benchmark, all method predicts a comparable number of contacts, around 240 for ordered proteins and around 45 for disordered ones, with Boltz-2 as the only outlier, predicting 50% more short-range (i,i+4) contacts for disordered proteins (Figure S2). All predictors also globally agree well on which contacts are present, but less so for disordered proteins: 84–93% of contacts are shared between predictor pairs for ordered proteins, versus 50–85% for disordered proteins (Figure S3A,B), where agreement is slightly lower for long-range contacts (Figure S3C,D). This trend extends to the full predicted distance distributions: for residue pairs with a contact predicted by at least one predictor, the two predictors’ distance distributions are more similar (lower mean Wasserstein distance) for ordered than for disordered proteins (Figure S4).

The distributions of average distances computed from the distograms are similar across predictors (Figure 1B), with disordered proteins showing a larger proportion of long distances and a peak at 1 nm that is proportionally smaller for ordered proteins. The average distances are well correlated between predictors, with mean pairwise Pearson coefficients of 0.76 for ordered and 0.71 for disordered proteins (Figure S5). The width of the distograms is more strongly affected by disorder: every predictor shows a cluster of high-spread, low-confidence predictions (standard deviation around 0.27 nm) for disordered proteins that is almost absent for ordered ones (Figure 1C). In contrast, the correlation of the distogram width between predictors is unaffacted by the presence of disorder (0.68 and 0.67), and sporadic predictor-specific differences persist at all disorder levels, suggesting that they reflect architecture and training rather than a disorder-dependent effect.

Overall, predicted distograms are broadly consistent across predictors, with greater protein-specific variability in the predictions for disordered proteins. In IDPs, stronger disagreements for longer-range contacts are observed in the benchmark, which is consistent with our analysis of PaaA2 distograms. This prediction variability further motivates the use of an explicit uncertainty model, along with a physics-based prior when generating ensembles that integrate AI structure predictors, as implemented in bAIes. Because distograms have no direct ground truth to be compared against, it remains unclear whether a specific approach is more accurate than the others in predicting the structural features of IDPs. To shed light on this point, we used bAIes to generate ensembles with different predictors and compared them with experiments, as presented in the following sections.

### Computational efficiency and scalability of bAIes

To optimize computational efficiency, we implemented bAIes natively in OpenMM.^44^ Since GPU acceleration is expected to benefit large proteins the most, we first benchmarked its computational efficiency across platforms and hardware as a function of protein size.

We first compared the LAMMPS^31,50^ and OpenMM implementations on our prior coil model as a function of protein size. To do this, we constructed disordered polyQ chains of 10 to 500 residues and measured simulation speed in ns/day (Figure 2A). As expected, the OpenMM GPU implementation becomes increasingly advantageous with chain length, reaching 379 ns/day for a 500-residue polyQ protein, compared to 200 ns/day for the LAMMPS CPU implementation at the same time step (2 fs). For chains shorter than about 100 residues, however, the LAMMPS CPU implementation was faster, with speeds decreasing from nearly 400 to 1000 ns/day over this range, compared to 1500 to 600 ns/day for OpenMM on GPU.

**Figure 2:**
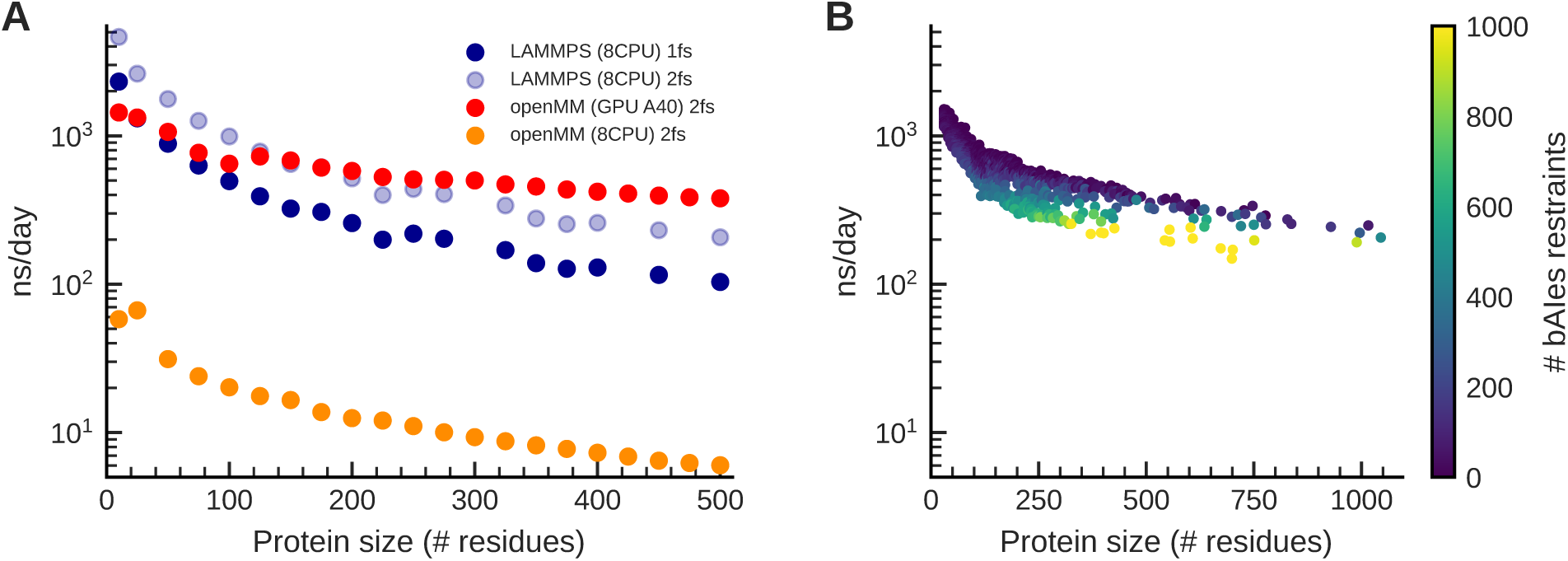
Efficient and scalable implementation of bAIes in OpenMM. **A**: Simulation speed (ns/day) of our coil model for polyQ chains using LAMMPS (blue) and OpenMM (orange and red) as a function of protein size. **B**: Simulation speed (ns/day) of bAIes using OpenMM as a function of protein size. Each point represents a single bAIes simulation of a disordered region from the DisProt database (n = 2883), performed on an NVIDIA A40 GPU with a 2 fs time step. Points are colored according to the number of pairwise bAIes restraints (see colorbar).

Next, we evaluated the performance of bAIes in OpenMM as a function of the number of bAIes restraints and protein size with simulations of 2883 disordered regions longer than 30 residues from the DisProt database.^52,65^ These regions ranged from 30 to 1045 residues and had between 0 and 2256 pairwise bAIes restraints per simulation. The results are summarized in Figure 2B. Simulation speed depended on protein size in the same way as for the coil models (Figure 2A). The overhead of the bAIes restraints was mostly noticeable for proteins below 200 residues and didn’t significantly slow down the computational cost of the simulations (Figure S6).

### Experimental validation of bAIes across AI structure predictors

We first evaluated bAIes with each of the six AI structure predictors on our previously published benchmark of 17 IDPs, which comprises experimental NMR and SAXS data^31^ (Table S1). This dataset is composed of three categories: IDPs with secondary structure motifs, for which bAIes is most relevant; fully disordered proteins with little structural information; and IDPs with prediction errors, on which bAIes has been previously shown to be particularly robust. For each category and predictor, we generated structural ensembles and back-calculated NMR and SAXS data. Figure 3A shows the *C_α_* chemical shift RMSE for each bAIes model for all IDPs with residual secondary structure motifs. For AF2, the new ensembles reproduce our previous finding:^31^ bAIes agrees better with experiments than the coil model. Overall, all structure predictors yield a similar improvement in agreement with chemical shifts relative to the coil model, suggesting improved local conformational sampling. For PaaA2, both local and global conformational sampling are improved, as indicated by the SAXS, RDC, and chemical shift profiles (Figure 3B-E, S7). For fully disordered proteins (Figure S8A) and IDPs with prediction errors (Figure S8B), all bAIes ensembles likewise agree equally well with experimental chemical shifts, confirming that the robustness of bAIes is transferable to the other structure predictors. Finally, the conditionally folded protein *Spm − FrpC* ^66^ is predicted to be folded by all predictors, highlighting a common limitation of these tools in capturing environment-dependent conformations.

**Figure 3:**
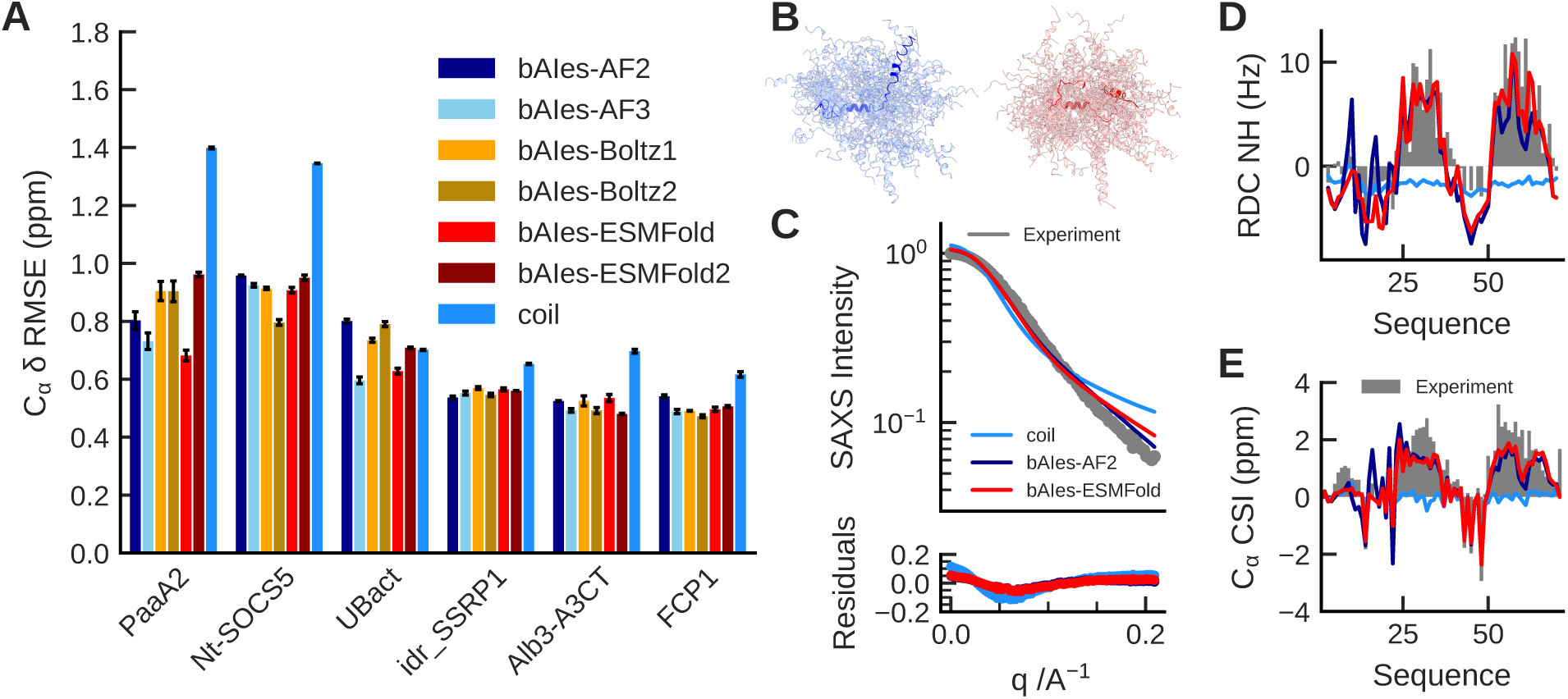
Validation of bAIes ensembles against experimental observables across structure predictors. **A**: RMSE of bAIes (AF2: dark blue, AF3: light blue, Boltz-1: yellow, Boltz-2: ochre, ESMFold: red, ESMFold2: dark red) and coil ensembles against experimental backbone *C_α_* chemical shifts for six partially ordered IDPs, calculated using SPARTA+. Error bars were computed with blocking analysis, in which the 1,500 simulation frames were divided into three independent blocks of 500 frames each (Methods). **B**: Structural ensembles of PaaA2 generated with bAIes-AF2 (blue) and bAIes-ESMFold (red). **C–E**: Experimental (grey) and calculated observables for PaaA2 using bAIes-AF2 (blue) and bAIes-ESMFold (red): **C**, SAXS profiles; **D**, NH residual dipolar couplings; and **E**, *C_α_* chemical shift index.

Next, we compared the accuracy of bAIes with atomistic MD simulations on the four proteins for which we have extensive NMR, SAXS, and all-atom simulation data: A*β*40, PaaA2, ACTR and *α*-synuclein. The bAIes ensembles built from all of the six predictors are overall competitive with all-atom simulations (Figure 4A and S9). In particular, the ESMFold-based bAIes ensembles agree with experiments better than, or within the error of, those from a99SB-disp, a state-of-the-art force field for IDPs.^15^ For *α*-synuclein and ACTR, the results are consistent across the bAIes ensembles, indicating that the six predictors provide similar structural information for these proteins. For A*β*40 and PaaA2, in contrast, accuracy varies between predictors, with AF2 and ESMFold giving the most accurate ensembles. For A*β*40, which behaves almost like a random coil in solution,^67^ predictors that yield more contacts resulted in bAIes ensembles with higher *RMSE_N_* (Figure S10). For PaaA2, relative accuracy depends mostly on the number of predicted long-range N-terminal contacts (Figure 1A). Many of those contacts (bAIes-AF2, bAIes-AF3) improved agreement with SAXS, whereas fewer improved agreement with RDCs and chemical shifts (Figures 1A,S9), suggesting that these contacts are transient or absent. ESMFold, which predicted few long-range contacts, yields a bAIes ensemble that agrees best overall, balancing local and global sampling accuracy.

**Figure 4:**
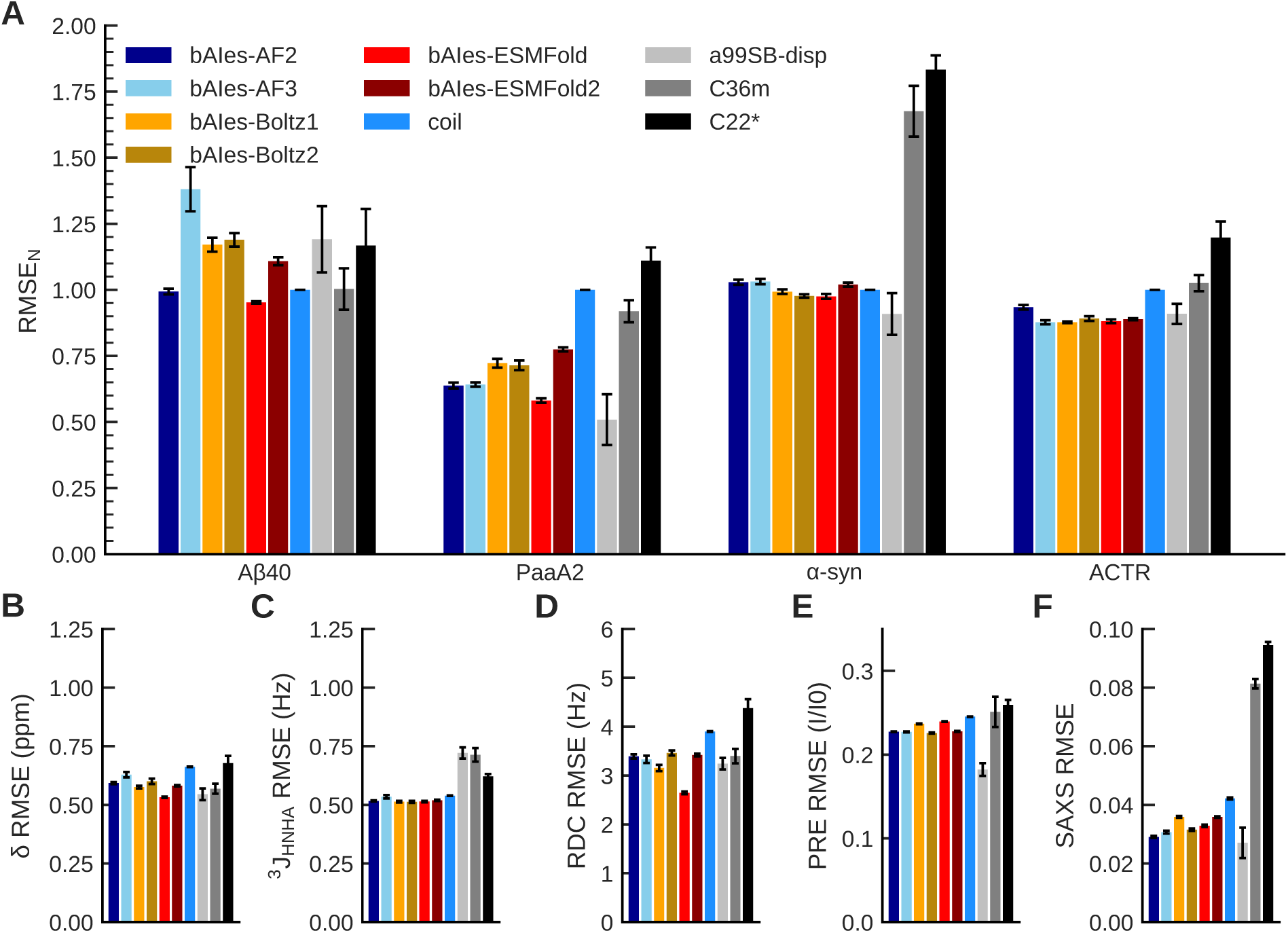
Accuracy of bAIes ensembles compared with state-of-the-art all-atom simulations. **A**: Normalized root mean square error (RMSE*_N_*) for A*β*40, PaaA2, *α*-synuclein, and ACTR, comparing bAIes ensembles generated using AF2 (dark blue), AF3 (light blue), Boltz-1 (yellow), Boltz-2 (ochre), ESMFold (red), and ESMFold2 (dark red); coil ensembles (blue); and all-atom simulations using amber99SB-disp (light grey), CHARMM36m (dark grey), and CHARMM22* (black). **B–F**: Average errors across the same four-protein benchmark for each experimental observable, with models shown in the same order and colors as in A: **B**, chemical shifts calculated using SPARTA+; **C**, ^3^*J*_HNHA_ couplings; **D**, RDCs; **E**, PRE intensity ratios; and **F**, SAXS profiles. Error bars were obtained from blocking analysis, in which the simulation frames were divided into three blocks containing an equal number of frames (Methods).

**Figure 5:**
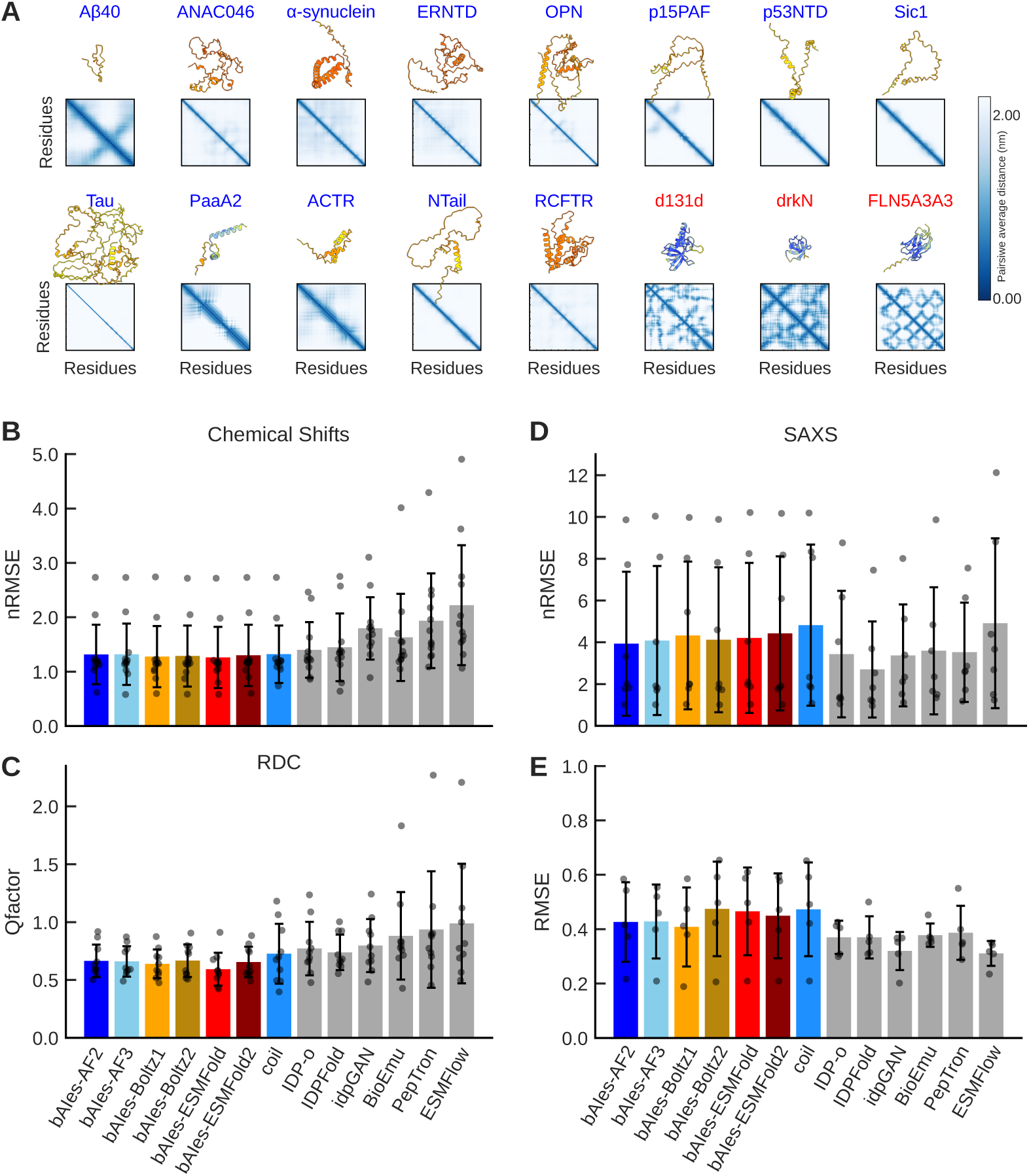
Benchmarking of bAIes against state-of-the-art IDP ensemble generation methods. **A**: ESMFold predictions for proteins in the PeptoneDB-Integrative benchmark, showing predicted structures (colored by pLDDT) and the corresponding pair-wise average distance matrices. **B**: Normalized RMSE (see Methods) between predicted and experimental chemical shifts (n = 12). Models compared include bAIes ensembles generated using AF2 (dark blue), AF3 (light blue), Boltz-1 (yellow), Boltz-2 (ochre), ESMFold (red), and ESMFold2 (dark red); coil ensembles (dodger blue); and six additional ensemble-generation methods (grey, from left to right: IDP-o, IDPFold, idpGAN, BioEmu, PepTron, and ESMFlow). **C**: Q-factors measuring agreement between predicted and experimental residual dipolar couplings (n = 10). **D**: Normalized RMSE between predicted and experimental SAXS profiles (n = 7). **E**: RMSE between predicted and experimental PRE intensity ratios (n = 5). Across all panels, models are shown in the same order and with the same colors. Bars represent the mean, error bars indicate the standard deviation across proteins, and grey markers show individual proteins. Each panel includes the subset of benchmark proteins for which the corresponding experimental data are available, excluding drkN, d131d, and FLN5A3A3 (main text and Figure S12).40

When the results are decomposed by experimental observable, bAIes best reproduces the local conformational sampling probed by chemical shifts, RDCs and ^3^J-couplings (Figure 4B-D). All structure predictors give satisfactory results, with the ESMFold-based ensembles slightly superior on these specific experimental data types. Agreement is weaker for all bAIes ensembles on PREs and SAXS, which report on long-range contacts and overall chain compaction, respectively (Figure 4E,F). This is consistent with the lower inter-predictor agreement on long-range contacts observed in this work (Figure S3C,D) and mirrors our earlier observations with AF2.^31^ It also fits with the predictors being trained on folded structures, whose training sets contain little information on dynamic and transient interactions.

### Benchmarking of bAIes against other IDP ensemble generation methods

To further test our method, we compared bAIes with other state-of-the-art IDP ensemble generators on the PeptoneDB-Integrative dataset^28^ (Table S2). The dataset contains 16 wellcharacterized IDPs with experimental NMR and SAXS data, four of which (A*β*40, PaaA2, ACTR and α-synuclein) were also in the previous benchmark against all-atom simulations. The dataset includes NMR chemical shifts, RDCs, PREs, and SAXS profiles, which together report on local conformational properties, long-range contacts, and global compaction of each IDP.

We first examined the properties of each protein and its structure predictions. The experimental chemical shifts (Figure S11) indicate that most proteins in this dataset are almost fully disordered, with few structural features (13 proteins with a g-score *>* 0.75), while a few, such as PaaA2, NTail, and ACTR, contain regions with helical propensity. Figure 5A shows the average distance matrix from the ESMFold distograms, together with the predicted ESMFold model of each protein. Most proteins appear disordered in the predictions, but three predictions (labeled in red in Figure 5A) are folded, with high mean pLDDT scores ranging from 70 to above 90 depending on the predictor (Table S3).

These three proteins belong to a class of systems with condition-dependent folding, which structure predictors tend to predict as folded, as for spm FrpC, which folds only with calcium ions^66^ and drkN (slow folded-disordered state exchange^31,68^). Δ131Δ (d131d), a low-stability staphylococcal nuclease fragment folding upon ligand binding,^69,70^ and FLN5A3A3, a protein variant designed to stay unfolded,^71^ are two additional examples. Since most structure predictor-dependent ensemble generators (bAIes, IDPFold, BioEmu, PepTron, ESMFlow) predict these proteins as folded, to avoid favoring ensemble generators that always produce disordered conformations (coil, IDP-o), we excluded them from the ensemble analysis. Results including them are reported in Supplementary Information (Figure S12).

For each protein, we generated bAIes ensembles with each of the six AI structure predictors, back-calculated the experimental observables, and compared them with experiments. We also compared our ensembles with those obtained with six other IDP ensemble generators: IDP-o,^28^ a fragment-based chain-growth method that assembles plausible IDP conformations from six-residue fragments drawn from the AlphaFold Protein Structure Database; ^72^ IDPFold,^29^ an SE(3) diffusion model pretrained on X-ray and NMR structures from the PDB and fine-tuned on atomistic conformations back-mapped from coarse-grained CALVADOS simulations of the IDRome; ^13^ idpGAN, a generative adversarial network trained on coarse-grained MD simulations of disordered sequences that generates C*α* coordinates from sequence;^25^ BioEmu, a diffusion model conditioned on AF2 Evoformer embeddings, trained on AlphaFold Database structures and extensive MD simulations and fine-tuned on experimental folding stabilities, which generates backbone-frame ensembles;^27^ ESMFlow, a flow-matching model that repurposes ESMFold as an ensemble generator and is trained on PDB structures;^30^ and PepTron, which shares the ESMFlow architecture but is additionally fine-tuned on synthetic ensembles of IDRome^13^ sequences generated with IDP-o, to better capture disordered regions.^28^

The benchmark results are shown in Figure 5B-E for each type of experimental data. On chemical shifts, the bAIes models are on average the most accurate, together with the coil model, followed closely by IDP-o and IDPFold (Figure 5B). The same trend holds for RDCs, with bAIes-ESMFold slightly ahead, followed by the other bAIes models, our coil model, IDPFold, IDP-o and idpGAN (Figure 5C). These results confirm that bAIes reproduces local conformational sampling as well as or better than other state-of-the-art ensemble generators. When the three proteins with folded predictions are included (Figure S12), our coil model leads the benchmark on these data, followed closely by IDP-o. In contrast, bAIes performs more modestly on SAXS and PREs, with larger standard deviations and poorer agreement with experiments than the other methods (Figure 5D,E). This is consistent with structure predictors producing few or no long-range contacts for disordered proteins. Taken together, these results indicate that bAIes matches state-of-the-art generators on local conformational properties, while the accuracy of long-range and global properties is limited by the information provided by the AI structure predictors.

## Discussions

In this work, we extended bAIes to six AI state-of-the-art structure predictors: AF2, AF3, Boltz-1, Boltz-2, ESMFold and ESMFold2. Comparing the predictors showed that IDP distograms are largely consistent, with sporadic disagreements that motivate an uncertainty-aware approach such as bAIes. In addition, bAIes was on average comparably accurate against experimental data using any of the predictors. This low sensitivity to the choice of predictor likely stems from the noise model of bAIes, which downweights unreliable restraints and, as we previously showed, makes the method robust to prediction errors.^31^ A few erroneous, low-confidence long-range contacts thus have little influence on the ensemble.

We benchmarked bAIes against all-atom MD simulations and state-of-the-art IDP ensemble generators. Although the total number of proteins is limited (29), they are well characterized, with several types of experimental data reporting on both local and global conformational properties. This allows for a clearer assessment of the strengths and weaknesses of each predicted ensemble relative to experiments. In all benchmarks, bAIes was strongest at reproducing data that report on local conformational sampling (chemical shifts, ^3^J-couplings, and RDCs), indicating that the local structural features in the distograms of each predictor are correctly transcribed into ensembles. On the mostly disordered PeptoneDB-Integrative set, bAIes outperforms the other models on these observables, closely followed by the coil prior. Thus, the added value of bAIes is clearest for proteins in presence of structured motifs and residual secondary structure.

In contrast, with every predictor, bAIes faced difficulties in reproducing the experimental observables sensitive to longe-range properties, namely chain compaction (SAXS) and long-range contacts (PREs). Whereas global observables such as SAXS profiles and average radii of gyration can be readily matched by reweighting, as we showed in our previous work,^31^ PRE data are considerably harder to reproduce even upon reweighting. Because PREs report on specific long-range contacts, the probability that an unbiased ensemble contains a sub-ensemble with the correct proportion of such contacts decreases exponentially with chain length. To alleviate this, we propose two solutions: (i) a more complex physico-chemical prior, with sufficiently accurate long-range interaction terms to reproduce PREs together with the radius of gyration and SAXS profiles. This is a promising direction for large-scale ensemble prediction; (ii) integration of experimental data during sampling, including PREs when available.^73,74^

A second limitation is shared by all predictors: proteins with condition-dependent folding (Spm-FrpC, drkN, d131d, and FLN5A3A3) are systematically predicted as folded, and bAIes inherits this behavior. The limitation applies to most ensemble generators that rely on the PDB or on structure predictors, which is why we excluded these proteins from the ensemble comparison. Users should therefore be cautious with proteins known or suspected to fold conditionally, since the AI-derived information does not reflect ligands, ions, or other environmental factors.

Given our results, no AI structure predictor stands out for use with bAIes. Although bAIes-ESMFold performed slightly better on local conformations, the size of the benchmark does not allow us to conclude that ESMFold systematically outperforms the other predictors; thus, all the tested predictors appear appropriate choices to perform bAIes simulations. We therefore suggest running several predictors and selecting either the most consensual prediction or, since consensus is not a safeguard in every case as we have seen for conditionally folded proteins, the one most compatible with the available experimental information. When computational resources are limited, ESMFold-based predictors are a suitable choice, both practically and conceptually: they are computationally efficient, yield satisfactory results with bAIes, and extract structural features from a protein language model without multiple sequence alignments, which are not always meaningful for often poorly conserved disordered regions.

Finally, excluding computationally prohibitive all-atom simulations, bAIes is distinctive among IDP ensemble generators in being MD-based and therefore inherently adaptable. The new OpenMM implementation makes it readily extensible, allowing users to add custom restraints, combine force fields, or include GAFF-parametrized molecules with only a few lines of Python, while remaining fast on GPU (379 ns/day for a 500-residue chain) with only a small overhead from the bAIes restraints. This flexibility makes bAIes compatible with a broad range of integrative modeling inputs, including experimental restraints and, as shown here, AI-derived information from any of the tested structure predictors. Many ML-based ensemble generators are limited by restricted input formats and by their inability to account for environmental conditions such as pH, temperature, or ionic strength, which MD-based approaches can, in principle, incorporate through the physical model and simulation conditions. We therefore expect bAIes to be particularly well suited to specific biological studies, in which AI-derived information can be combined with (i) system-adapted physico-chemical priors and (ii) experimental data. In the future, we will pursue these two directions to improve the accuracy of bAIes and make it a practical and versatile tool for integrative modeling of disordered proteins.

## Acknowledgments

This project has received funding from the European Research Council (ERC) under the European Union’s Horizon 2020 research and innovation programme (Grant agreement No. 101086685 – bAIes). The authors would like to thank Julian Olivier Streit for useful discussions.

## Author contributions

M.B. and V.S. conceived and designed the research. V.S., N.P., and M.B. wrote, revised and edited the manuscript. V.S designed, performed, and analyzed the simulations. V.S. developed the OpenMM bAIes implementation. N.P. modified the AI structure predictors to provide distograms.

