## Supplementary Information for "Leveraging state-of-the-art AI structure predictors to generate atomic resolution ensembles of intrinsically disordered proteins"

#### Supporting Information

Vincent Schnapka,* Nicolas Portal, and Massimiliano Bonomi*

*Institut Pasteur, Université Paris Cité, CNRS UMR 3528, Computational Structural Biology Unit, Paris, France*

*

### Supplementary Figures

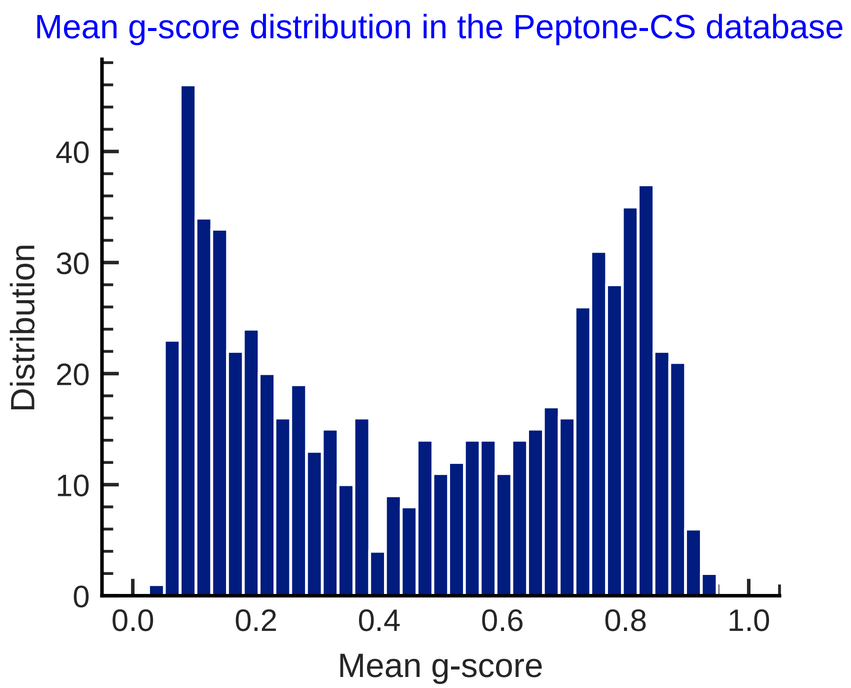


**Supplementary Figure 1: Distribution of disorder g-scores in the PeptoneDB-CS dataset.** Distribution of mean per-protein g-scores across the 659 proteins in PeptoneDB-CS, comprising 329 ordered proteins (g-score $< 0.5$) and 330 disordered proteins (g-score $> 0.5$).

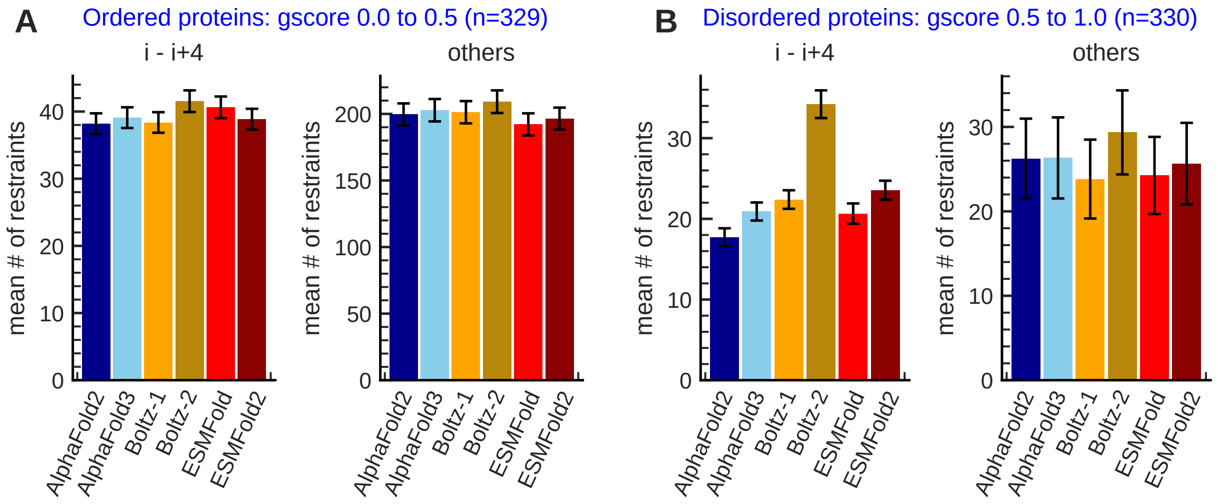


**Supplementary Figure 2: Comparison of predicted inter-residue contacts across structure predictors in PeptoneDB-CS.** Mean number of predicted inter-residue contacts per protein for **A**, ordered proteins (mean g-score $< 0.5$, n $= 329$), and **B**, disordered proteins (mean g-score $> 0.5$, n $= 330$) from the PeptoneDB-CS dataset. For each group, contacts are separated into short-range contacts between residues separated by four positions in the sequence ($i, i+4$; left) and all other contacts (right). Predictions are shown for AlphaFold2 (dark blue), AlphaFold3 (light blue), Boltz-1 (yellow), Boltz-2 (ochre), ESMFold (red), and ESMFold2 (dark red). Error bars were obtained using bootstrap analysis.

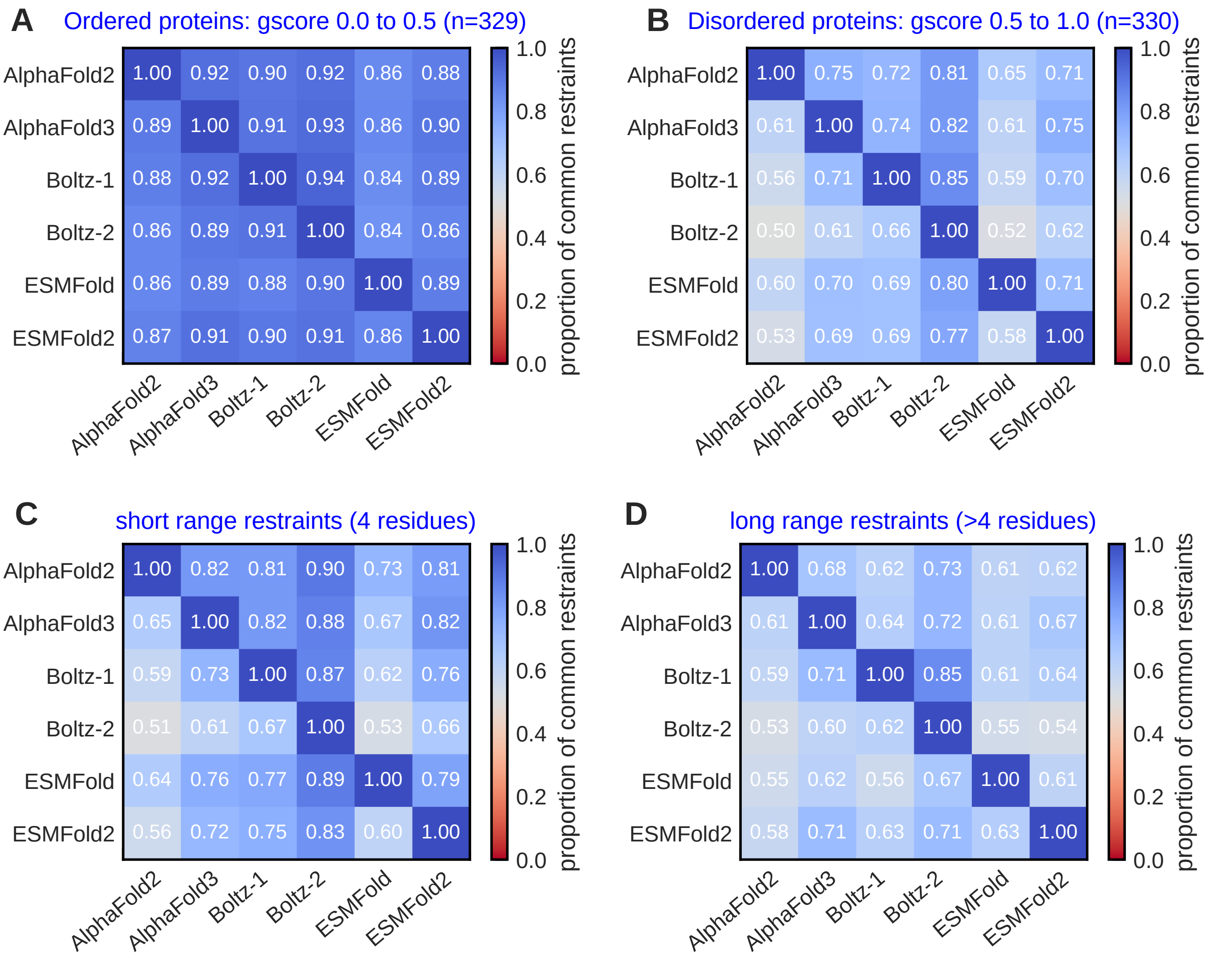


**Supplementary Figure 3: Agreement between structure predictors on inter-residue contacts in PeptoneDB-CS.** Pairwise consensus matrices showing the proportion of predicted inter-residue contacts shared between the six structure predictors (Methods). Rows and columns correspond to AlphaFold2, AlphaFold3, Boltz-1, Boltz-2, ESMFold, and ESMFold2. **A**: Ordered proteins (mean g-score $< 0.5$, n $= 329$). **B**: Disordered proteins (mean g-score $> 0.5$, n $= 330$). **C**: Short-range contacts between residues separated by four positions in the sequence $(i, i+4)$ in disordered proteins. **D**: Longer-range contacts between residues separated by more than four positions in the sequence in disordered proteins.

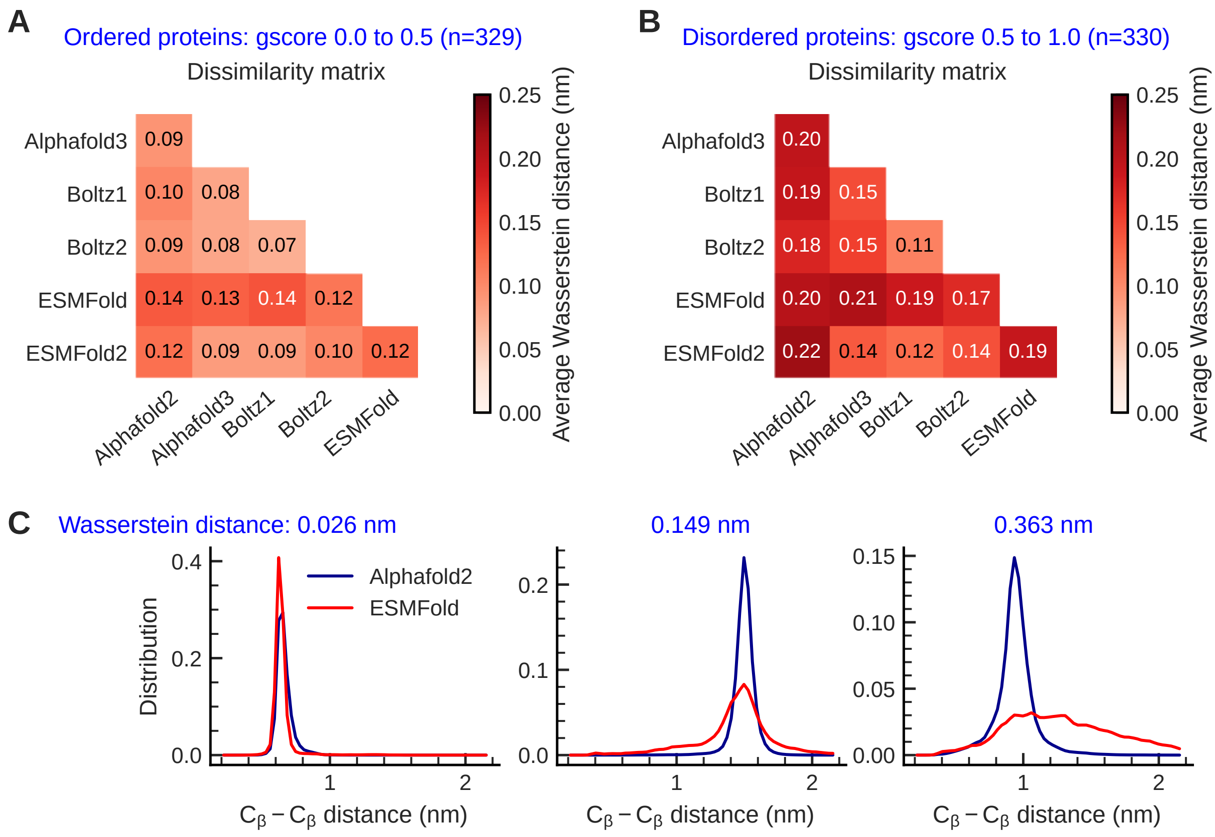


**Supplementary Figure 4: Comparison of predicted distograms across structure predictors using Wasserstein distances.** **A–B**: Matrices of average pairwise Wasserstein distances between the six structure predictors for **A**, ordered proteins (mean g-score $< 0.5$, n $= 329$), and **B**, disordered proteins (mean g-score $> 0.5$, n $= 330$), from the PeptoneDB-CS dataset. Wasserstein distances were calculated between predicted distograms for residue pairs associated with at least one inter-residue contact among the six predictors, then averaged over residue pairs and proteins. Larger values indicate greater differences between the predicted distance distributions. **C**: Representative comparisons of individual distograms predicted by AlphaFold2 (blue) and ESMFold (red), illustrating increasing Wasserstein distances between the two distributions (0.026, 0.149, and 0.363 nm, from left to right).

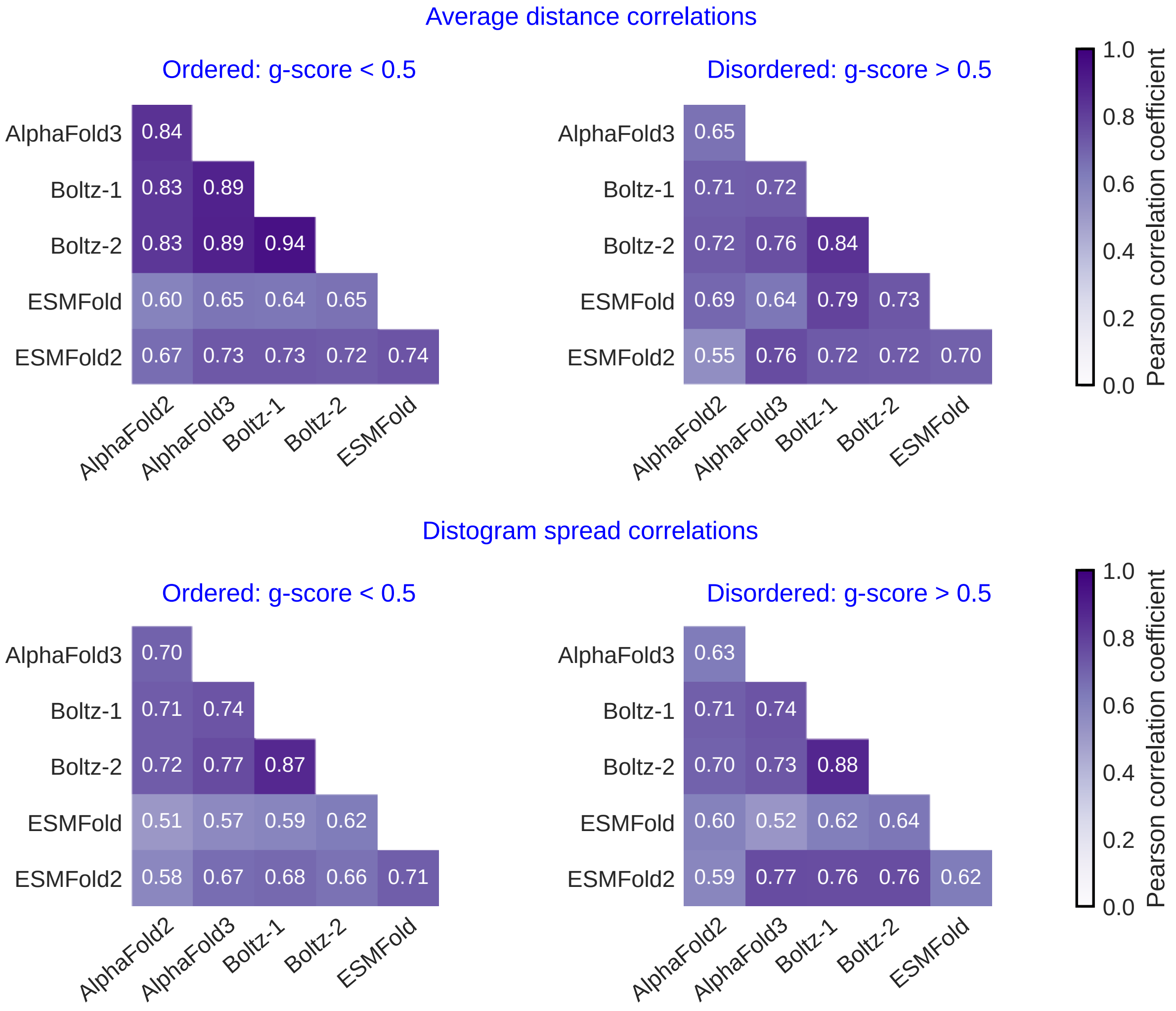


**Supplementary Figure 5: Correlations of predicted distogram mean distances and spreads across structure predictors.** Pairwise Pearson correlation matrices comparing the six structure predictors for ordered (left, mean g-score $< 0.5$, n $= 329$) and disordered proteins (right, mean g-score $> 0.5$, n $= 330$) from the PeptoneDB-CS dataset. The top row shows correlations between distogram mean distances, while the bottom row shows correlations between distogram standard deviations (spreads). For each protein and pair of predictors, Pearson correlation coefficients were calculated using the mean distances or standard deviations of residue pairs associated with at least one predicted inter-residue contact among the six predictors.

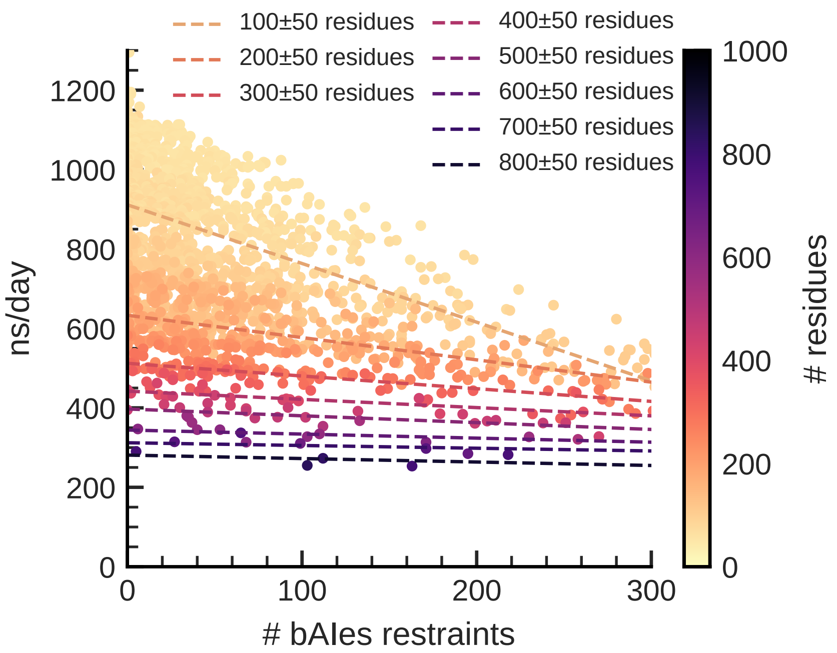


**Supplementary Figure 6: Dependence of bAIes simulation speed on the number of restraints and protein size.** Simulation speed (ns/day) of bAIes in OpenMM as a function of the number of pairwise bAIes restraints, evaluated on disordered regions from the DisProt database. Each point represents a single bAIes simulation, colored according to protein size (see colorbar). Dashed lines represent linear fits to subsets of simulations grouped into eight protein-size windows, ranging from 150–250 to 750–850 residues, in increments of 100 residues.

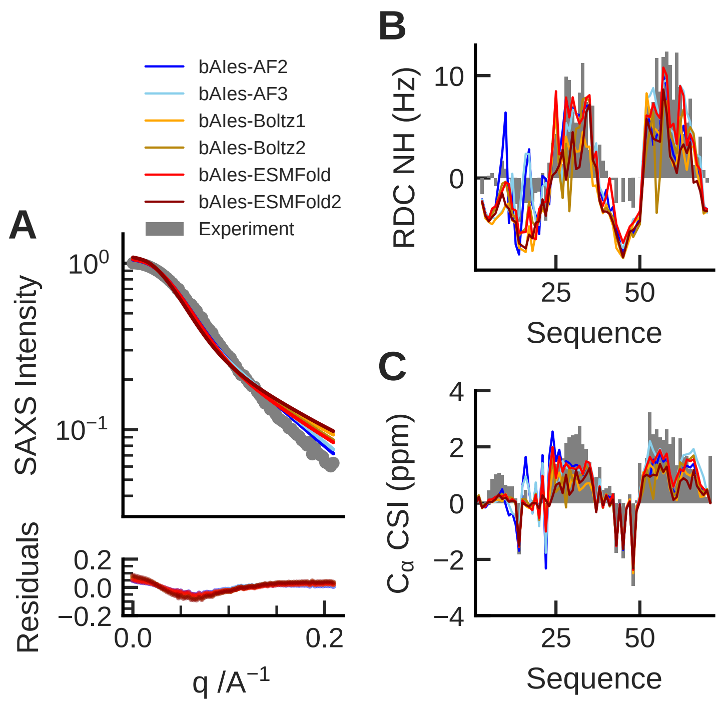


**Supplementary Figure 7: Agreement of PaaA2 bAIes ensembles with experimental SAXS, RDC, and chemical shift data.** Comparison of experimental measurements (grey) with observables calculated from PaaA2 ensembles generated using bAIes with six structure predictors: AlphaFold2 (dark blue), AlphaFold3 (light blue), Boltz-1 (yellow), Boltz-2 (ochre), ESMFold (red), and ESMFold2 (dark red). **A**: Small-angle X-ray scattering (SAXS) intensity profiles, with residuals shown below. **B**: Residual dipolar coupling (RDC) profiles as a function of residue position. **C**: $C_\alpha$ chemical shift indices (CSI) as a function of residue position.

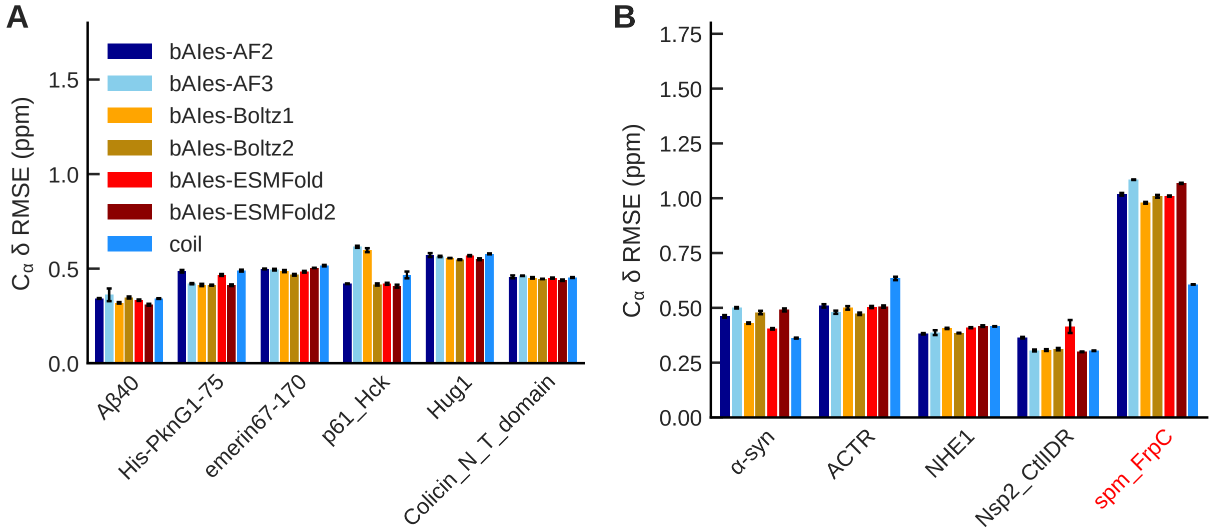


**Supplementary Figure 8: Agreement of bAIes ensembles with experimental chemical shifts across IDP classes.** Average root-mean-square errors (RMSEs) between calculated and experimental $C_\alpha$ chemical shifts for **A**, six fully disordered IDPs, and **B**, five IDPs with structure prediction errors, from the 17-IDP benchmark. Results are shown for bAIes ensembles generated using AlphaFold2 (dark blue), AlphaFold3 (light blue), Boltz-1 (yellow), Boltz-2 (ochre), ESMFold (red), and ESMFold2 (dark red), as well as random-coil ensembles (dodger blue). Error bars were obtained by blocking analysis, dividing the 1,500 simulation frames into three independent blocks of 500 frames each (see Methods).

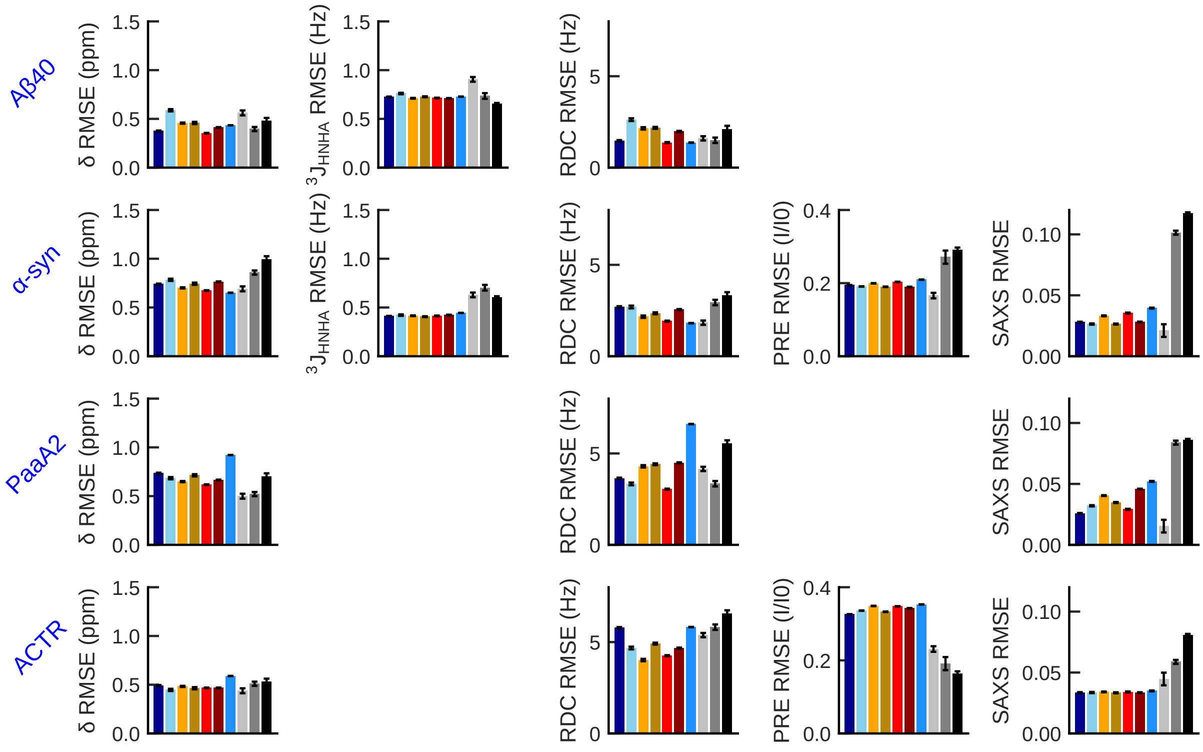


**Supplementary Figure 9: Agreement of bAIes and all-atom MD ensembles with individual experimental observables.** Average root-mean-square errors (RMSEs) between calculated and experimental observables, including chemical shifts, J-couplings, residual dipolar couplings (RDCs), paramagnetic relaxation enhancements (PREs), and small-angle X-ray scattering (SAXS) profiles, for four IDPs with available all-atom simulation data: A$\beta$40, $\alpha$-synuclein, PaaA2, and ACTR. Each row shows the agreement with the experimental data types available for the corresponding protein. Results are shown for bAIes ensembles generated using AlphaFold2 (dark blue), AlphaFold3 (light blue), Boltz-1 (yellow), Boltz-2 (ochre), ESMFold (red), and ESMFold2 (dark red), random-coil ensembles (dodger blue), and ensembles generated using three state-of-the-art all-atom MD force fields: a99SB-disp (light grey), CHARMM36m (grey), and CHARMM22* (black). Error bars were obtained by blocking analysis, dividing the simulation frames into three independent blocks containing equal numbers of frames (Methods).

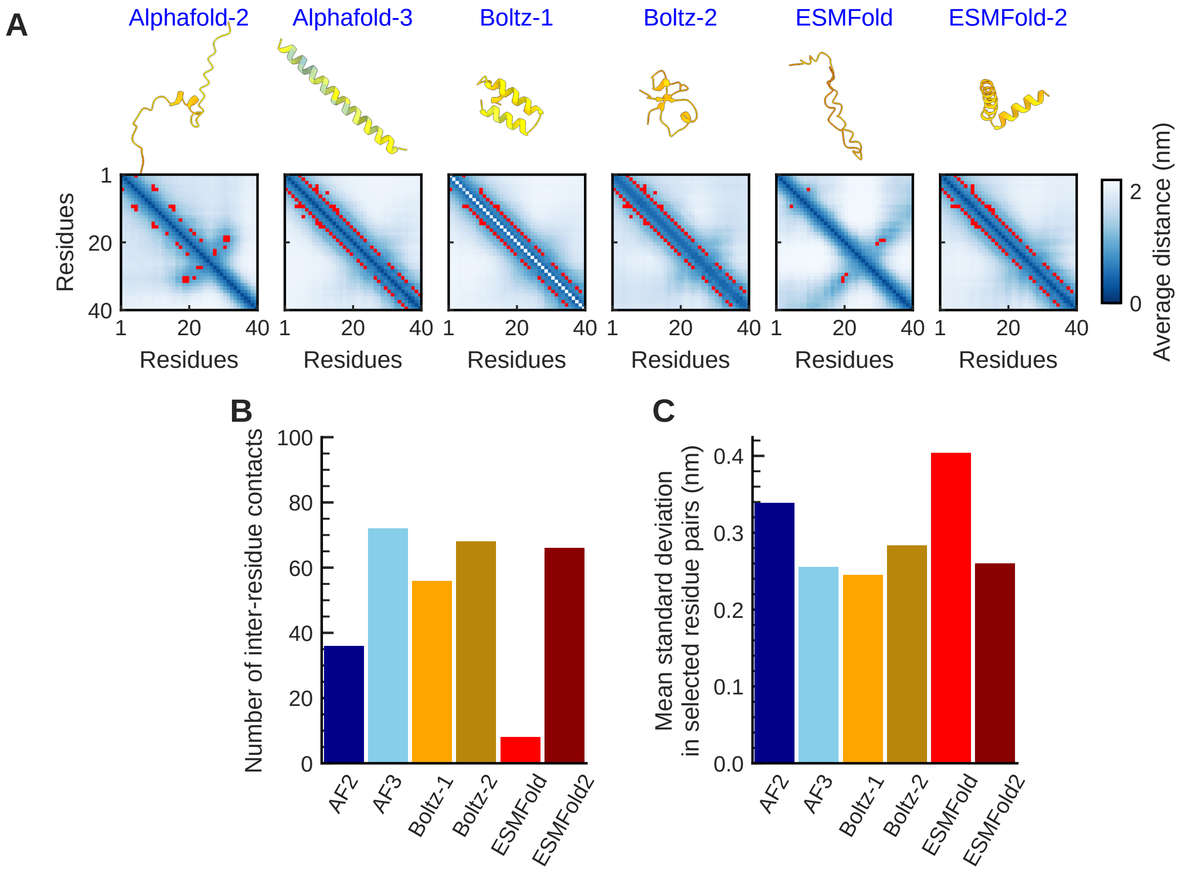


**Supplementary Figure 10: Comparison of A$\beta$40 structure predictions and distogram-derived restraints across predictors.** **A**: Predicted structures and corresponding distogram mean-distance matrices for A$\beta$40 across the six structure predictors: AlphaFold2, AlphaFold3, Boltz-1, Boltz-2, ESMFold, and ESMFold2. Predicted structures are colored according to their per-residue pLDDT scores, while residue pairs identified as inter-residue contacts and used as restraints in bAIes are highlighted in red in the matrices. **B**: Number of predicted inter-residue contacts (bAIes restraints) for each structure predictor. **C**: Mean standard deviation of the predicted distance distributions across residue pairs selected as bAIes restraints for each predictor.

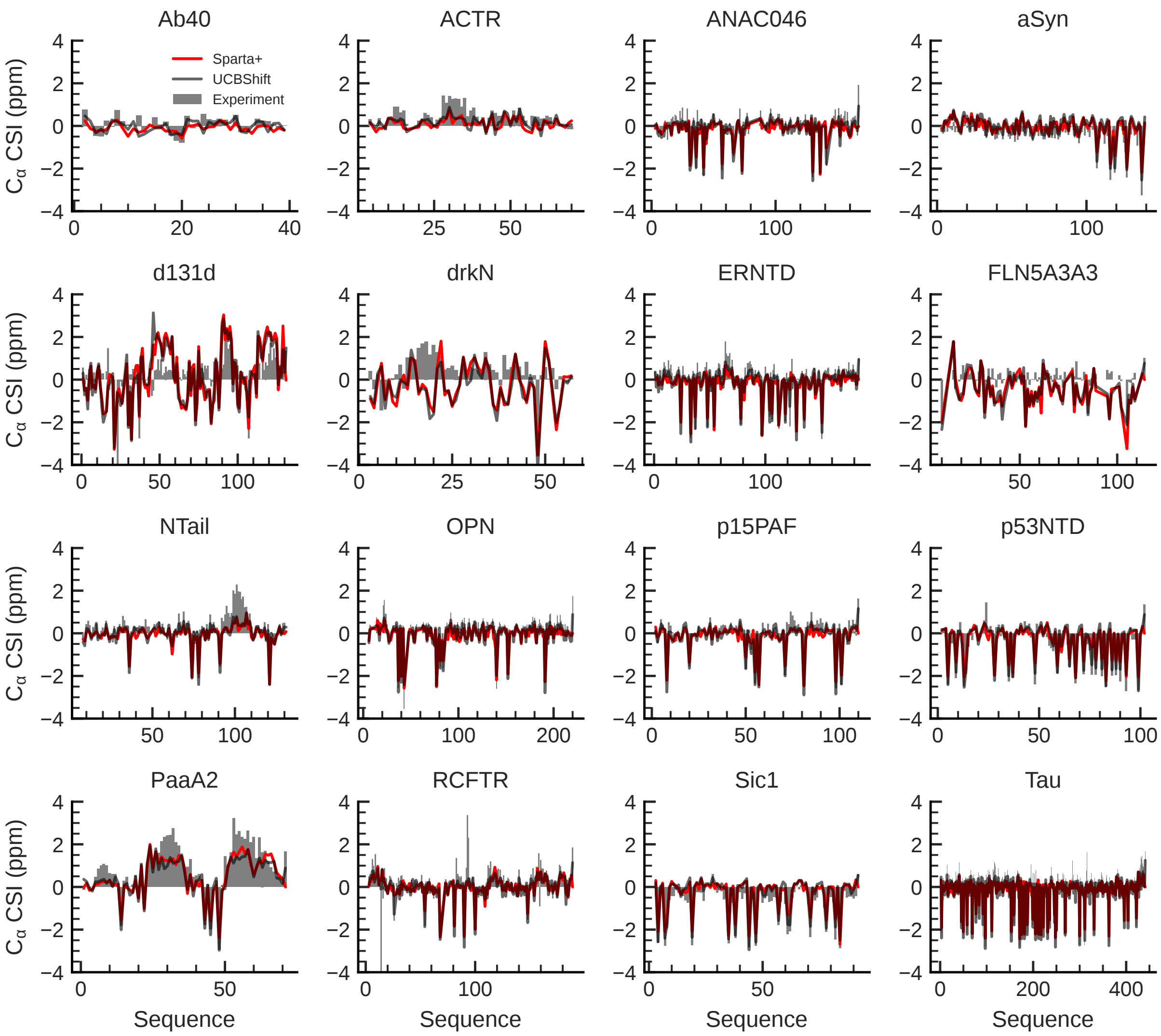


**Supplementary Figure 11: Comparison of experimental and predicted $C_\alpha$ secondary chemical shifts for PeptoneDB-Integrative IDPs.** Experimental $C_\alpha$ secondary chemical shifts (grey bars) and corresponding values back-calculated from bAIes-ESMFold ensembles using SPARTA+ (red) and UCBShift (black) for the 16 intrinsically disordered proteins (IDPs) in the PeptoneDB-Integrative dataset.

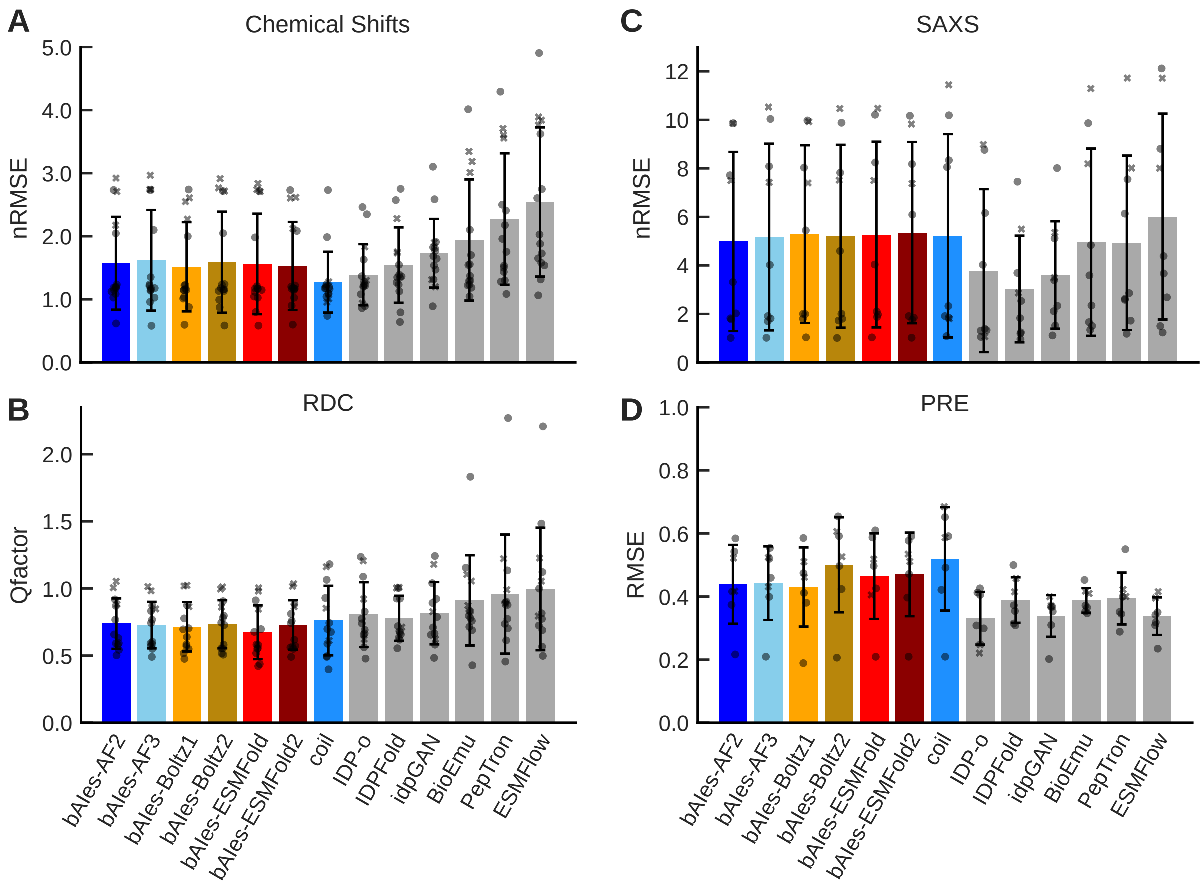


**Supplementary Figure 12: Benchmarking of bAIes against other IDP ensemble generation methods across all PeptoneDB-Integrative proteins.** Comparison of the agreement between experimental observables and ensembles generated using bAIes and other methods for the 16 IDPs in the PeptoneDB-Integrative dataset. **A**: Normalized root-mean-square errors (RMSEs; Methods) between calculated and experimental chemical shifts for 15 IDPs. **B**: Q-factors measuring agreement between calculated and experimental residual dipolar couplings (RDCs) for 13 IDPs. **C**: Normalized RMSEs between calculated and experimental small-angle X-ray scattering (SAXS) profiles for 9 IDPs. **D**: RMSEs between calculated and experimental paramagnetic relaxation enhancement (PRE) intensity ratios for 7 IDPs. All panels compare bAIes ensembles generated using AlphaFold2 (dark blue), AlphaFold3 (light blue), Boltz-1 (yellow), Boltz-2 (ochre), ESMFold (red), and ESMFold2 (dark red); random-coil ensembles (dodger blue); and six other ensemble generation methods (grey, from left to right: IDP-o, IDPFold, idpGAN, BioEmu, PepTron, and ESMFlow). Models are shown in the same order and colors across panels. Bars represent the mean across proteins, error bars indicate the standard deviation, and individual proteins are shown as grey markers. The three conditionally folded proteins, drkN, d131d, and FLN5A3A3, are highlighted with crosses. Each panel includes only proteins for which the corresponding experimental data are available.

### Supplementary Tables

**Supplementary Table 1: Summary of ensemble generation and all-atom simulations used to validate and benchmark bAIes.** List of simulations performed or analyzed for the 17-IDP benchmark dataset from Ref.,[1] used to evaluate bAIes ensembles generated with different structure predictors and compare them with all-atom molecular dynamics simulations. For each protein (row) and ensemble generation method (column), the reported integer indicates the number of conformers generated for random-coil and bAIes ensembles or analyzed for previously published all-atom simulations.

| **system** | coil, bAIes-AF2, -AF3, -Boltz-1, -Boltz-2, -ESMFold, -ESMFold2 | a99SB-disp[2] C36m,[3] C22*[4] | experimental data |
|---|---|---|---|
| **A$\beta$40** | 1500 | 30000 | CS, RDC, $^3$J |
| **PaaA2** | 1500 | 30000 | CS, RDC, SAXS |
| **$\alpha$-synuclein** | 1500 | 30000 | CS, RDC, PRE, SAXS, $^3$J |
| **ACTR** | 1500 | 30000 | CS, RDC, PRE, SAXS |
| **emerin67-170** | 1500 | - | CS |
| **p61_Hck** | 1500 | - | CS |
| **His-PknG1-75** | 1500 | - | CS |
| **Colicin_NT_domain** | 1500 | - | CS |
| **Hug1** | 1500 | - | CS |
| **Nt-SOCS5** | 1500 | - | CS |
| **UBact** | 1500 | - | CS |
| **idr_SSRP1** | 1500 | - | CS |
| **Alb3-A3CT** | 1500 | - | CS |
| **FCP1** | 1500 | - | CS |
| **NHE1** | 1500 | - | CS |
| **Nsp2_CtlIDR** | 1500 | - | CS |
| **spm_FrpC** | 1500 | - | CS |

**Supplementary Table 2: Summary of ensemble generation and simulations used to benchmark bAIes on PeptoneDB-Integrative.** List of simulations performed (first column) or analyzed (second column, ensembles taken from Invernizzi *et al.*[5]) for the 16 IDPs in the PeptoneDB-Integrative dataset, used to benchmark bAIes ensembles generated with different structure predictors against other protein ensemble generation methods. For each protein (row) and method (column), the reported integer indicates the number of conformers generated for random-coil and bAIes ensembles or analyzed for ensembles produced by the other methods.

| **system** | coil, bAIes-AF2, -AF3, -Boltz-1, -Boltz-2, -ESMFold, -ESMFold2 | IDP-o,[5] IDPFold,[6] idpGAN,[7] BioEmu,[8] PepTron,[5] ESMFlow[9] | experimental data[5] |
|---|---|---|---|
| **A$\beta$40** | 1500 | 1000 | CS, RDC |
| **ANAC046** | 1500 | 1000 | CS, SAXS, PRE |
| **$\alpha$-synuclein** | 1500 | 1000 | CS, RDC, SAXS, PRE |
| **ERNTD** | 1500 | 1000 | CS, SAXS, PRE |
| **OPN** | 1500 | 1000 | CS, RDC, PRE |
| **p15PAF** | 1500 | 1000 | CS, RDC, SAXS |
| **p53NTD** | 1500 | 1000 | CS, RDC |
| **Sic1** | 1500 | 1000 | CS, RDC, SAXS, PRE |
| **Tau** | 1500 | 1000 | CS, RDC, SAXS, PRE |
| **PaaA2** | 1500 | 1000 | CS, RDC, SAXS |
| **ACTR** | 1500 | 1000 | CS, RDC, SAXS, PRE |
| **NTail** | 1500 | 1000 | CS, RDC |
| **RCFTR** | 1500 | 1000 | CS, RDC |
| **$\Delta$131$\Delta$** | 1500 | 1000 | CS, RDC |
| **drkN** | 1500 | 1000 | CS, RDC, SAXS, PRE |
| **FLN5A3A3** | 1500 | 1000 | CS, RDC, SAXS, PRE |

**Supplementary Table 3: Comparison of mean pLDDT scores across structure predictors in PeptoneDB-Integrative.** Mean predicted local distance difference test (pLDDT) scores for each protein and structure predictor in the 16-IDP PeptoneDB-Integrative benchmark. Scores above 80 are highlighted, as are proteins for which structure predictors systematically predict folded conformations.

| prediction | af2 | af3 | boltz1 | boltz2 | esm | esm2 |
|---|---|---|---|---|---|---|
| Ab40 | 56.3 | 81.7 | 64.2 | 50.9 | 47.7 | 54.6 |
| ANAC046 | 53.3 | 59.8 | 43.6 | 33.8 | 34.3 | 76.2 |
| aSyn | 65.5 | 78.5 | 54.3 | 48.2 | 33.2 | 73.6 |
| ERNTD | 54.5 | 57.5 | 51.5 | 34.5 | 31.9 | 71.2 |
| OPN | 56.8 | 61.7 | 48.8 | 36.2 | 43.9 | 76.1 |
| p15PAF | 65.5 | 69.2 | 56.8 | 44.4 | 46.7 | 48.1 |
| p53NTD | 62.3 | 67.8 | 58.4 | 51.3 | 54.5 | 50.9 |
| Sic1 | 65.1 | 60.1 | 52.6 | 44.7 | 51.2 | 42.7 |
| Tau | 50.4 | 57.5 | 43.9 | 30.6 | 57.1 | 78.5 |
| PaaA2 | **91.8** | **88.7** | 66.3 | **86.8** | 76.8 | 61.8 |
| ACTR | 70.9 | 79.9 | 62.3 | 60.9 | 49.9 | 54.7 |
| NTail | 65.7 | 61.8 | 56.6 | 42.1 | 48.2 | 43.4 |
| RCFTR | 59.4 | 63.3 | 52.7 | 41.9 | 35.8 | 73.9 |
| **d131d** | **95.6** | **96.9** | **93.6** | **94.8** | **93.8** | **81.5** |
| **drkN** | **87.7** | **96.1** | **82.5** | **94.9** | **91.8** | 75.2 |
| **FLN5A3A3** | **91.6** | **93.3** | **83.5** | **88.7** | **88.8** | 72.1 |
